# Posttranslational modification of the RHO of plants protein RACB by phosphorylation and cross-kingdom conserved ubiquitination

**DOI:** 10.1101/2020.05.28.121228

**Authors:** Lukas Weiß, Lana Gaelings, Tina Reiner, Julia Mergner, Bernhard Kuster, Attila Fehér, Götz Hensel, Manfred Gahrtz, Jochen Kumlehn, Stefan Engelhardt, Ralph Hückelhoven

## Abstract

Small RHO-type G-proteins act as signaling hubs and master regulators of polarity in eukaryotic cells. Their activity is tightly controlled, as defective RHO signaling leads to aberrant growth and developmental defects. Two major processes regulate G-protein activity: canonical shuttling between different nucleotide bound states and posttranslational modification (PTM), of which the latter can support or suppress RHO signaling, depending on the individual PTM. In plants, regulation of Rho of plants (ROPs) signaling activity has been shown to act through nucleotide exchange and GTP hydrolysis, as well as through lipid modification, but there is little data available on phosphorylation or ubiquitination of ROPs. Hence, we applied proteomic analyses to identify PTMs of the barley ROP RACB. We observed *in vitro* phosphorylation by barley ROP binding kinase 1 and *in vivo* ubiquitination of RACB. Comparative analyses of the newly identified RACB phosphosites and human RHO protein phosphosites revealed conservation of modified amino acid residues, but no overlap of actual phosphorylation patterns. However, the identified RACB ubiquitination site is conserved in all ROPs from *Hordeum vulgare*, *Arabidopsis thaliana* and *Oryza sativa* and in mammalian Rac1 and Rac3. Point mutation of this ubiquitination site leads to stabilization of RACB. Hence, this highly conserved lysine residue may regulate protein stability across different kingdoms.

## Introduction

Small guanosine triphosphate-binding (G-) proteins are master regulators of developmental and immunity-related processes including membrane transport, nuclear protein trafficking, reactive oxygen species production and cytoskeleton organization in growth and immunity (Irani et al., 1997; Lin and Yang, 1997; Jones et al., 2002; Yang, 2008; Ebine et al., 2011; Akamatsu et al., 2013). In humans (*Homo sapiens*, *Hs*), small G-proteins are organized in the rat sarcoma (RAS) superfamily, comprised of the ARF, RAB, RAN, RAS and RHO families (Wennerberg et al., 2005), with the latter being further divided into Cdc42-like, Rho-like, RAC-like, RhoBTB and Rnd subfamilies (Lawson and Ridley, 2018). Plants possess ARF, RAB and RAN family proteins, but only one type of small G-proteins from the RAS homologue (RHO) family, so called Rho of plants (ROPs). Traditionally, they were also called plant RACs (RAS-related C3 botulinum toxin substrate), due to their slightly higher similarity to metazoan RACs than RHO proteins. ROPs can be further distinguished based on their C-terminal hypervariable region and associated lipid modification. Type I ROPs can be reversibly *S*-acylated and additionally contain a CaaX-box motif that is a target for constitutive cysteine prenylation. Type II ROPs lack the CaaX-motif and are apparently constitutively *S*-acylated. Those cysteine lipid modifications are essential for plasma membrane anchoring and downstream signaling (Sorek et al., 2011; Yalovsky, 2015). Additionally, RHO proteins are often called molecular switches, because they cycle between a guanosine diphosphate (GDP)-bound “OFF” state and the signaling-competent guanosine triphosphate (GTP)-bound “ON” state. This cycling itself is tightly controlled by specific proteins: guanosine nucleotide exchange factors (GEFs) facilitate the GDP to GTP exchange, which leads to G-protein activation. In turn, GTPase activating proteins (GAPs) target GTP-bound G-proteins and stimulate their weak intrinsic GTPase activity, which results in GTP hydrolysis and turns the G-proteins signaling incompetent (Berken and Wittinghofer, 2008). This nucleotide exchange is accompanied by a conformational shift, which affects the interaction with other proteins (Milburn et al., 1990; Berken, 2006). A third class of regulatory proteins, so-called guanine nucleotide dissociation inhibitors (GDIs), targets mostly the GDP-bound state and recruits the G-protein to the cytoplasm, blocking it from downstream signaling (Boulter and Garcia-Mata, 2010).

Lipidations are not the only posttranslational modifications (PTM) in RHO proteins. In animals, both phosphorylation and ubiquitination are employed to control RHO signaling, thereby enhancing or suppressing G-protein function in a case-dependent manner. The mammalian RHO family member Rac1 for instance is phosphorylated at three amino acids and ubiquitinated at two residues (reviewed in Abdrabou and Wang (2018)), whereas for mammalian RhoA four phosphorylation sites and three ubiquitination sites have been identified (Lang et al., 1996; Forget et al., 2002; Ellerbroek et al., 2003; Ozdamar et al., 2005; Rolli-Derkinderen et al., 2005; Wei et al., 2013; Tang et al., 2014; Tong et al., 2016) (see S1 Fig for an overview of Rac1 and RhoA PTMs).

Compared to the mammalian field, knowledge about phosphorylation and ubiquitination in ROPs is limited. To date, *in vitro* phosphorylation of *Arabidopsis thaliana* (At) ROP4 and ROP6 by ROP binding protein kinase 1 (RBK1), a receptor-like cytoplasmic kinase (RLCK) of class VI_A, has been reported. RBK1 further interacts with Mitogen-activated protein kinase 1 (MPK1) of an auxin-responsive MPK cascade, and it has been suggested that regulation of ROP4 and ROP6 via phosphorylation might be important for auxin-dependent cell expansion (Enders et al., 2017). Furthermore, type II AtROP10 and AtROP11 are phosphorylated at S138, but no functional studies about these phosphosites have been conducted yet (Mergner et al., 2020). Introduction of a phosphomimetic mutation of the broadly conserved S74 in AtROP4 and *Medicago sativa* MsROP6 has been investigated. This S74E mutation did not alter GTPase activity, but blocked interaction with an upstream activating ROPGEF, weakened the binding to a downstream signaling kinase and stimulated less *in vitro* kinase activity (Fodor-Dunai et al., 2011). However, there is no reported evidence for *in vivo* phosphorylation of type I ROPs.

ROPs have been also investigated in the interaction between barley (*Hordeum vulgare, Hv*) and the powdery mildew fungus *Blumeria graminis* f.sp. *hordei* (*Bgh*). There it was shown that most barley ROPs can support the penetration success of *Bgh* (Schultheiss et al., 2002; Schultheiss et al., 2003; Pathuri et al., 2008; Hoefle et al., 2011). Among these barley ROPs, RACB is the best characterized ROP family member. Under physiological conditions, it is required for root hair outgrowth and stomatal subsidiary cell formation, and RACB and its interacting proteins further steer cytoskeleton organization (Engelhardt et al., 2019). During *Bgh* attack, RNA-interference (RNAi)-mediated knockdown of RACB decreased the penetration success of the fungus into barley epidermal cells, whereas overexpression of a constitutively activated RACB variant (CA, RACB-G15V, a GTPase mutant that cannot hydrolyse GTP) promoted susceptibility (Schultheiss et al., 2002; Schultheiss et al., 2003; Pathuri et al., 2008; Hoefle et al., 2011). In contrast, transient overexpression of neither wildtype (WT) nor a dominant negative RACB form (DN, RACB-T20N, a mutant that can only bind GDP) increased barley susceptibility, indicating that the GTP-bound, signaling-active form of RACB supports fungal invasion into intact barley leaf epidermal cells.

Since RHO family proteins interact with several downstream executors to perform their signaling function, interaction partners of RACB were investigated with regard to barley susceptibility to *Bgh*. It was shown that two classes of scaffolding proteins, Interactor of Constitutively Active ROPs/ROP-interactive partners (ICRs/RIPs) and ROP-interactive and CRIB-motif containing proteins (RICs), associate with CA-RACB at the plasma membrane and enhance penetration success of *Bgh* into barley epidermal cells when transiently overexpressed (Schultheiss et al., 2008; Engelhardt et al., 2019; McCollum et al., 2020). In contrast, two other interactors of activated RACB support resistance to fungal penetration. Microtubule-Associated ROP GTPase Activating Protein 1 (MAGAP1) enhances resistance towards *Bgh* and likely negatively regulates RACB by supporting GTP hydrolysis (Hoefle et al., 2011). Barley RBK1 interacts with CA-RACB in yeast and *in planta*, and CA-RACB increases RBK1 kinase activity *in vitro*. RBK1 may be a RACB-regulated kinase, since co-expression of CA-RACB and RBK1 results in co-localization of both proteins at the plant plasma membrane via recruitment of the otherwise cytosolic RBK1. However, unlike RACB, transient knockdown of RBK1 results in increased disease susceptibility and reduced microtubule stability (Huesmann et al., 2012). S-phase kinase 1-associated protein-like (SKP1L), an interactor of RBK1, is also involved in resistance towards *Bgh* (Reiner et al., 2016). SKP1L is a predicted subunit of an SCF E3 ubiquitin ligase complex, which targets specific proteins for polyubiquitination and proteasomal degradation (Hua and Vierstra, 2011). In line with this predicted function, it was demonstrated that RBK1, SKP1L and the 26S proteasome negatively influence protein abundance of transiently overexpressed CA-RACB. These proteins hence presumably act in a negative feedback-loop involving phosphorylation and ubiquitination of RACB (Reiner et al., 2016).

In this study, we confirm that the kinase activity of RBK1 is substantially enhanced in the presence of CA-RACB, leading to strong autophosphorylation of RBK1 and transphosphorylation of CA-RACB *in vitro*. Phosphosite mapping revealed novel phosphorylated residues in RACB that were not previously described to be modified in other RHO family proteins. We further found that CA-RACB displays an ubiquitination-typical appearance of high molecular weight derivatives after immunoprecipitation. Mass spectrometry revealed that CA-RACB is ubiquitinated at K167. Mutational studies support that RACB K167 is involved in regulating protein stability, but otherwise did not interfere with RACB’s ability to recruit interaction partners or influence susceptibility of barley to *Bgh*. However, data demonstrate *in planta* ROP ubiquitination at a residue that is conserved in mammalian Rac1, Rac3 and all other ROPs of barley, rice and Arabidopsis.

## Results

### CA-RACB is phosphorylated by RBK1 *in vitro*

Since previous research suggested that RACB might be negatively regulated through the protein kinase RBK1 and ubiquitination (Reiner et al., 2016), we aimed to discover putative posttranslational modifications in RACB. First, we investigated the potential of RACB to be phosphorylated by RBK1, because it was previously shown that constitutively activated (CA-) RACB interacts with RBK1 *in vivo* and enhances RBK1’s kinase activity *in vitro* (Huesmann et al., 2012). Therefore, we carried out an *in vitro* radiolabeling kinase assay with recombinantly expressed and purified RBK1 and CA-RACB, both C-terminally His-tagged (Fig 1A). C-terminal tagging should prevent artificial phosphorylation events, as Dorjgotov et al. (2009) showed RBK1-like RLCK VI_A kinases phosphorylating the N-terminal His-tag and linker-region of *Medicago sativa Ms*ROP6, but not the ROP protein itself. Since RBK1 alone shows some constitutive kinase activity and autophosphorylation *in vitro* (Huesmann et al., 2012), we used low RBK1 protein amounts in this assay to be able to see stimulated kinase activity. Under these conditions, only weak autophosphorylation of RBK1 was visible. In the presence of CA-RACB, however, we observed stronger autophosphorylation of RBK1 and transphosphorylation of CA-RACB. No phosphorylation could be detected after incubation with a RBK1 kinase-dead mutant (KD, K245D) or in a water-control (see Fig 1B for RBK1 protein stability).

**Fig 1.**
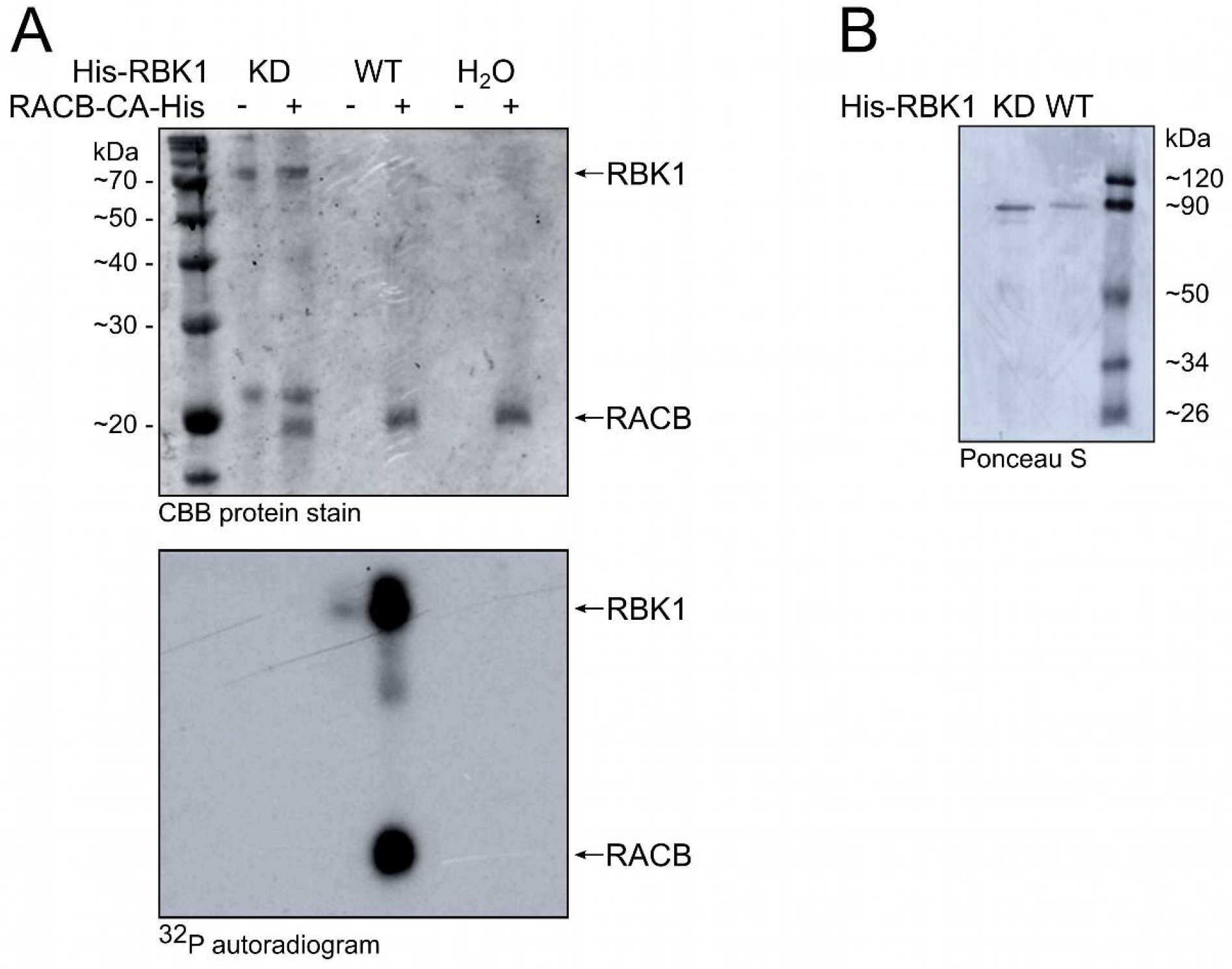
*In vitro* co-incubation of CA-RACB and RBK1 results in phosphorylation of both proteins. **(A)** Low amounts of His-tagged RBK1 alone showed weak autophosphorylation *in vitro*. Kinase activity was enhanced by the presence of constitutively activated His-tagged CA-RACB, resulting in extensive phosphorylation of both proteins. Upper panel = protein loading control. Lower panel = autoradiogram showing ^32^P-labelled, phosphorylated proteins. WT = wildtype RBK1; KD = kinase-dead RBK1 loss-of-function mutant (K245D); H_2_O = no kinase control. Estimations of molecular weight are depicted in kilo Dalton (kDa) on the left. **(B)** Ponceau S stain showing stability of purified WT and KD His-RBK1.

To eventually prove phosphorylation within the RACB protein sequence and to identify phosphosites in RACB, we conducted the *in vitro* kinase assay using RBK1 and unlabelled ATP and detected *in vitro* phosphosites of CA-RACB via mass spectrometry after tryptic digestion (Fig 2A). This revealed high confidence CA-RACB peptides with phosphorylated residues at positions S97, T139, S159, S160 and T162. Since in the case of the neighboring serines S159 and S160 the localization of the phospho-modification could not be specifically assigned within the peptide sequence (S1 Table), both residues were treated as possible but alternative phosphosites. Using computational modelling of the RACB protein structure based on the protein models of GDP-bound *At*ROP4 and GTP-analog-bound *Hs*RAC1, we suggest that S97, T139 and T162 are surface-exposed in RACB, whereas this is less clear for S159 and S160 (Fig 2B).

**Fig 2.**
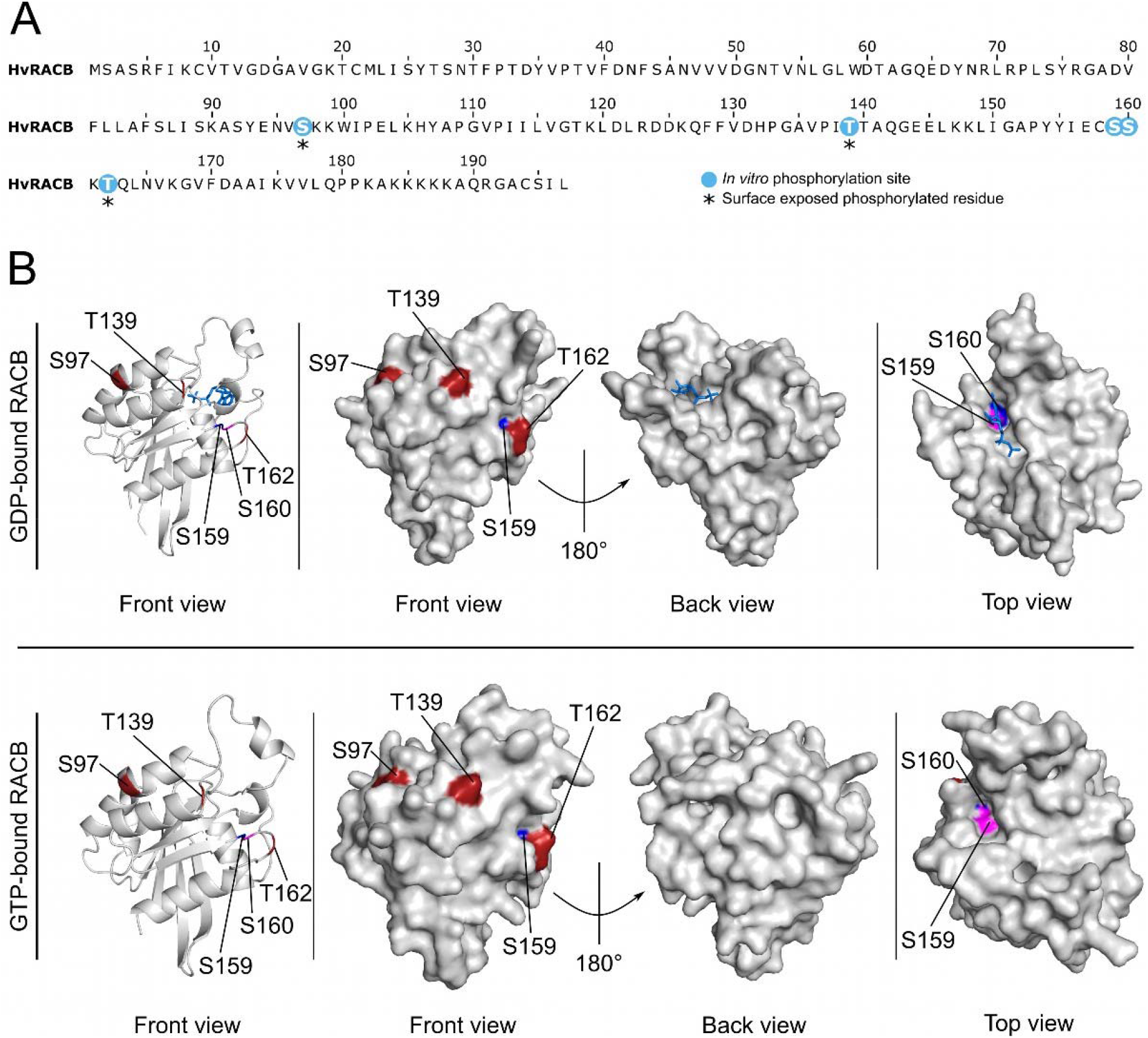
Surface-exposure of CA-RACB *in vitro* phosphosites. **(A)** Phosphorylation sites of CA-RACB are shown in blue or, when phosphorylated and predicted as surface-exposed, residues are additionally highlighted by asterisks. *In vitro* phosphorylation sites of RACB were identified by mass spectrometry after *in vitro* co-incubation of CA-RACB and RBK1. **(B)** Upper panel: the model of GDP-bound WT RACB (amino acids 1-178 from 197 total) was built with GDP-bound *At*ROP4 as template (PDB entry: 2NTY.1 (Thomas et al., 2007)). Lower panel: GTP-bound WT RACB (amino acids 7-178 from 197 total) was modelled using *Hs*RAC1 associated with GNP (phosphoaminophosphonic acid-guanylate ester, a non-hydrolysable GTP-analog) as template (PDB entry: 3TH5.1 (Krauthammer et al., 2012)). Phosphorylated surface-exposed residues are highlighted in red, GDP as ligand in blue. Only S97, T139 and T162 are surface-exposed, S159 (pink) and S160 (blue) are not clearly exposed to the surface. Putative RACB folds were assembled by homology-modelling using SWISS-MODEL and achieved GMQE (global model quality estimate) scores of 0.75 for GDP-RACB and 0.66 for GTP-RACB, and QMEAN DisCo (qualitative model energy analysis with consensus-based distance constraint) (Studer et al., 2020)) scores of 0.82 ± 0.07 and 0.80 ± 0.07, respectively. Surface structure (front, back and top view) and cartoon (front view) illustrations were modelled in PyMOL (Schrodinger, 2015).

Next, we tried to map *in vivo* phosphorylation sites for RACB. In order to obtain sufficient amounts of protein, we generated transgenic barley lines overexpressing HA-tagged CA-RACB, a cytosolic non-prenylated CA-RACB-ΔCSIL mutant (Schultheiss et al., 2003) or the HA-tag alone. Initially, we generated 47 transgenic plants for HA-CA-RACB, 37 plants for HA-RACB-ΔCSIL and 40 plants for HA alone. We propagated these lines and selected the offspring using PCR, resistance against the selection marker hygromycin and Western blotting (not shown). When we confirmed the expression of the HA-tagged RACB variants by Western blot, only the free HA-peptide could not be detected, most likely because of its small size (Fig 3A). Since CA-RACB is a susceptibility factor in the barley-*Bgh* interaction (Schultheiss et al., 2002; Schultheiss et al., 2003; Pathuri et al., 2008; Hoefle et al., 2011), we also scored the penetration efficiency of *Bgh* in leaf epidermal cells of all three transgenic lines to rule out any artificial effects of the HA-tag on the susceptibility function of RACB during the barley-*Bgh* interaction. Upon infection with *Bgh*, plants overexpressing full-length HA-CA-RACB displayed increased disease susceptibility compared to the free HA-control similar as shown before for untagged CA-RACB (Pathuri et al., 2008), whereas the HA-CA-RACB-ΔCSIL overexpressing plants did not show an increased disease phenotype (Fig 3B). This suggests that HA-tagged CA-RACB is functional in support of fungal penetration.

**Fig 3.**
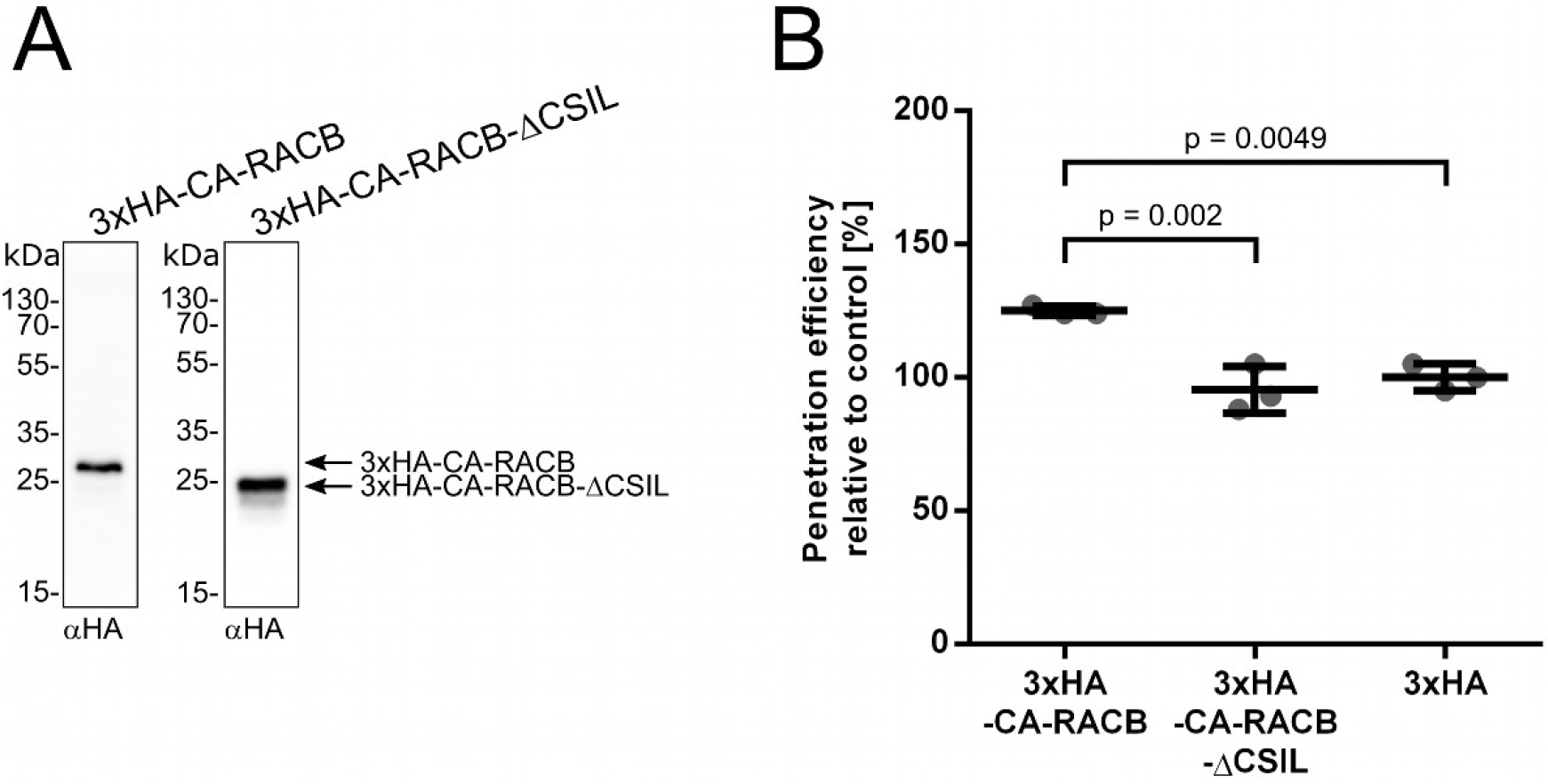
Transgenic barley plants overexpressing HA-tagged CA-RACB show increased *Bgh* susceptibility. **(A)** Both HA-tagged full-length CA-RACB and CA-RACB-ΔCSIL are stably expressed in transgenic barley. Eluate fractions of anti-HA immunoprecipitations were analyzed by anti-HA Western blotting. Plant material belonged to pools of transgenic lines BG657 E3 and E4 for *3xHA-CA-RACB* and BG658 E4 and E9 for *3xHA-CA-RACB*-ΔCSIL. **(B)** Leaf epidermal cells of transgenic plants expressing full-length HA-tagged CA-RACB were more susceptible towards penetration by *Bgh*, while plants expressing the truncated CA-RACB-ΔCSIL mutant show the same level of susceptibility as the HA-control. For each construct, three plants of one transgenic line were tested. Lines were BG657 E4 for 3x*HA*-CA-*RACB*, BG658 E4 for 3x*HA*-CA-*RACB*-ΔCSIL and BG659 E2 for 3x*HA*. Each datapoint corresponds to one plant with about 100 interaction sites scored, and means with standard deviation are shown. Statistical significances were determined using a One-way ANOVA with Tukey’s HSD.

To find *in vivo* phosphorylation sites of RACB, we initially isolated leaf mesophyll protoplasts from HA-CA-RACB overexpressing plants and, to eliminate any negative impact on RACB stability caused by phosphorylation as previously hypothesized (Reiner et al., 2016), we treated them with the proteasome inhibitor MG132. We extracted total proteins from these protoplasts and enriched HA-tagged CA-RACB using immunoprecipitation. Following tryptic digestion, we selected for phosphorylated peptides using Immobilized Metal Affinity Chromatography (IMAC). Peptides and posttranslational modifications were identified via mass spectrometry, but potential HA-CA-RACB phosphosites eluded our detection (not shown). Next, before MG132 treatment, we co-overexpressed GFP-tagged RBK1 in mesophyll protoplasts prepared from transgenic HA-CA-RACB overexpressing and control plants, and selectively skipped immunoprecipitation or IMAC (S2A Fig). This attempt also did not lead to the identification of *in vivo* RACB phosphopeptides, although more than 2,000 barley proteins were found to phosphorylated in those samples (S2 Table). In a third approach, we tried to map RACB phosphorylation after transient co-overexpression of GFP-RBK1 and proteasome inhibition by MG132 in transgenic HA-CA-RACB overexpressing barley mesophyll protoplasts. From this, we extracted proteins, performed a tryptic digest and enriched phosphopeptides via IMAC. These samples were subsequently decomplexed by high pH reversed-phase fractionation to increase the discovery rate of peptides during mass spectrometry. This identified more than 5,000 phosphopeptides from diverse barley proteins (S3 Table), thus providing a valuable resource for the barley research community. For instance, data show in vivo phosphorylation sites of ROP signaling components such as three different barley ROPGAPs (S4 Table). Because our experimental design included *in vivo* expression of combined or single GFP-tagged RBK1 and HA-CA-RACB (S2B Fig), some of the mapped barley phosphosites might be CA-RACB- and/or RBK1-dependent, but this would need further validation. Despite the high amount of identified phosphopeptides, no *in vivo* phosphopeptides of HA-CA-RACB were recovered.

Although we primarily aimed at identifying phosphopeptides of RACB, our datasets showed phosphosites for RBK1 (Fig 4). In the *in vitro* kinase assay, we found RBK1 to be phosphorylated at positions S26, S30, S98, S100, S107, T110, S155, T170 and S436. In the two experiments involving *in vivo* co-overexpression of CA-RACB and RBK1 (S2A Fig and Fig 4C), we found RBK1 phosphosites at positions S26, S30, S107, S113, S160, T170, S189, S197, S518, T521 and S26, S30, S100, S107, S113, S114, S216, T525, respectively. Overlapping sites from *in vitro* and at least one *in vivo* experiment are therefore S26, S30, S100, S107 and T170.

**Fig 4.**
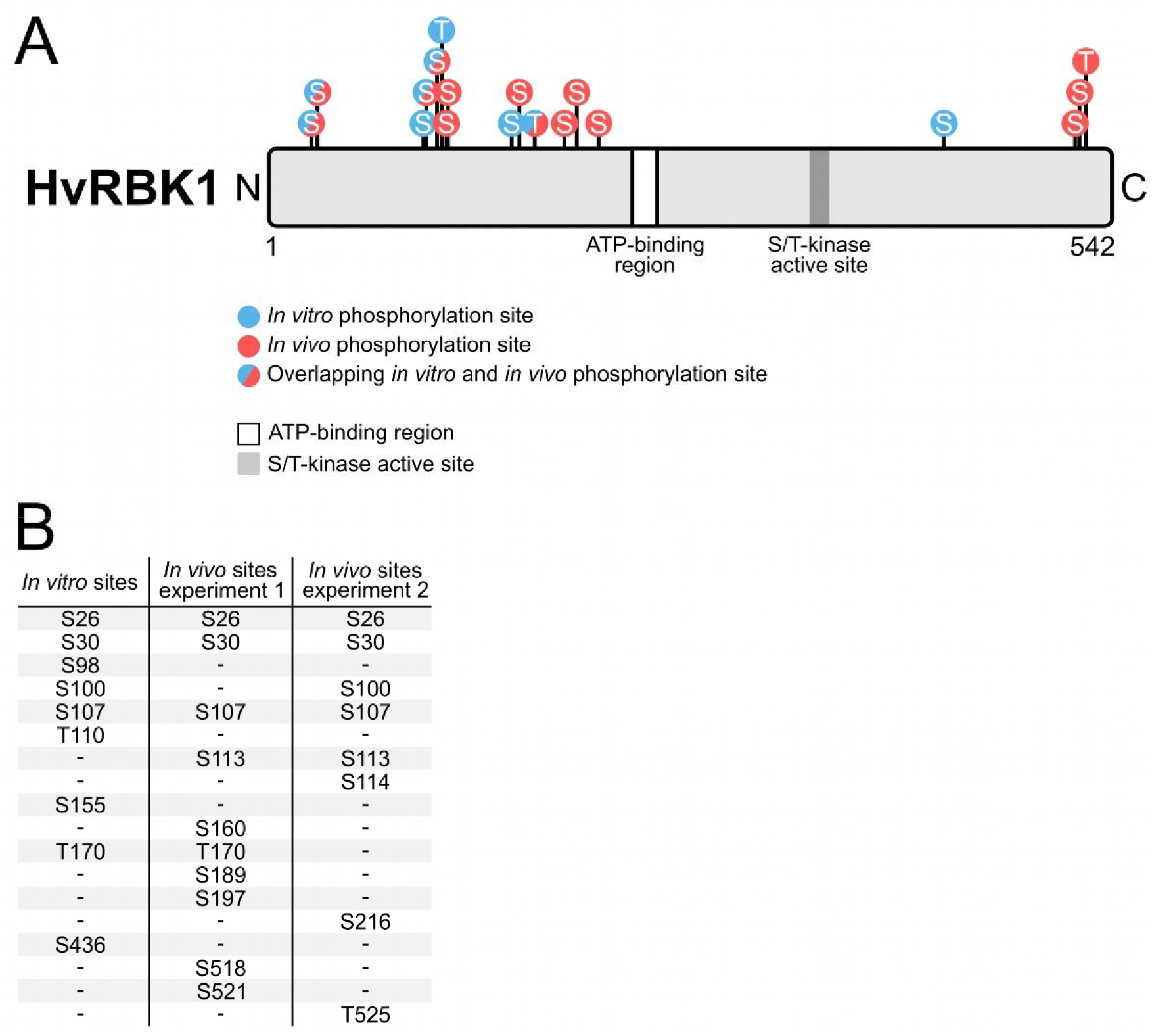
Comparison between *in vitro* and *in vivo* RBK1 phosphosites. **(A)** Schematic illustration of the barley RBK1 sequence with marked *in vitro* (blue) and *in vivo* (red) phosphosites. Residues phosphorylated under both conditions are shown with both colors. *In vitro* phosphorylated amino acids were identified in an *in vitro* kinase assay, in which His-tagged RBK1 and His-tagged CA-RACB were co-incubated. *In vivo* phosphosites were found in transgenic HA-CA-RACB overexpressing mesophyll protoplasts in the presence of transiently co-overexpressed GFP-tagged RBK1 and the proteasome-inhibitor MG132. Phosphopeptides were enriched from total protein extracts of protoplast lysates using IMAC (experiment 1) and additionally decomplexed by high pH reversed-phase fractionation (experiment 2). All *in vitro* and *in vivo* phosphosites were identified by mass spectrometry. **(B)** List of RBK1 phosphosites from all *in vitro* and *in vivo* experiments.

### CA-RACB is ubiquitinated *in vivo*

Previously, we found that the protein abundance of CA-RACB is negatively influenced by RBK1, SKP1L and a proteasome (Reiner et al., 2016). This prompted us to investigate potential *in vivo* CA-RACB ubiquitination. We used MG132 for proteasome inhibition in first leaves of transgenic barley overexpressing HA-tagged CA-RACB. This led to detection of HA-CA-RACB and higher molecular weight derivatives on Western blots after αHA-immunoprecipitation and detection via anti-HA antibodies (Fig 5A). To determine whether this band pattern is MG132 dependent, we repeated this experiment with a DMSO control and found a similar laddering pattern in the control (S3 Fig). As this laddering is usually a characteristic indicator for ubiquitination of a target protein (Gohre et al., 2008; Lu et al., 2011), we attempted to map the residue(s) at which HA-CA-RACB might be ubiquitinated. We first conducted a global mass spectrometry analysis, in which we specifically looked for branched amino acid sequences that derive from ubiquitinated lysine residues after tryptic digest (S2C Fig). Interestingly, we found a peptide containing RACB K167 carrying a mass shift equal to 2 glycine molecules (Fig 5B). This mass shift indicates that K167 is indeed ubiquitinated, as two glycine residues typically remain at ubiquitinated lysine residues (glyglycylation), whereas the rest of the ubiquitin moiety is cleaved by trypsin during sample digestion (Peng et al., 2003). Evidence for RACB ubiquitination was subsequently substantiated in a targeted parallel reaction monitoring (PRM) approach. Comparison with other ROPs from barley, rice and Arabidopsis showed that RACB K167 is a highly conserved residue in ROPs (Fig 6 and S4 Fig).

**Fig 5.**
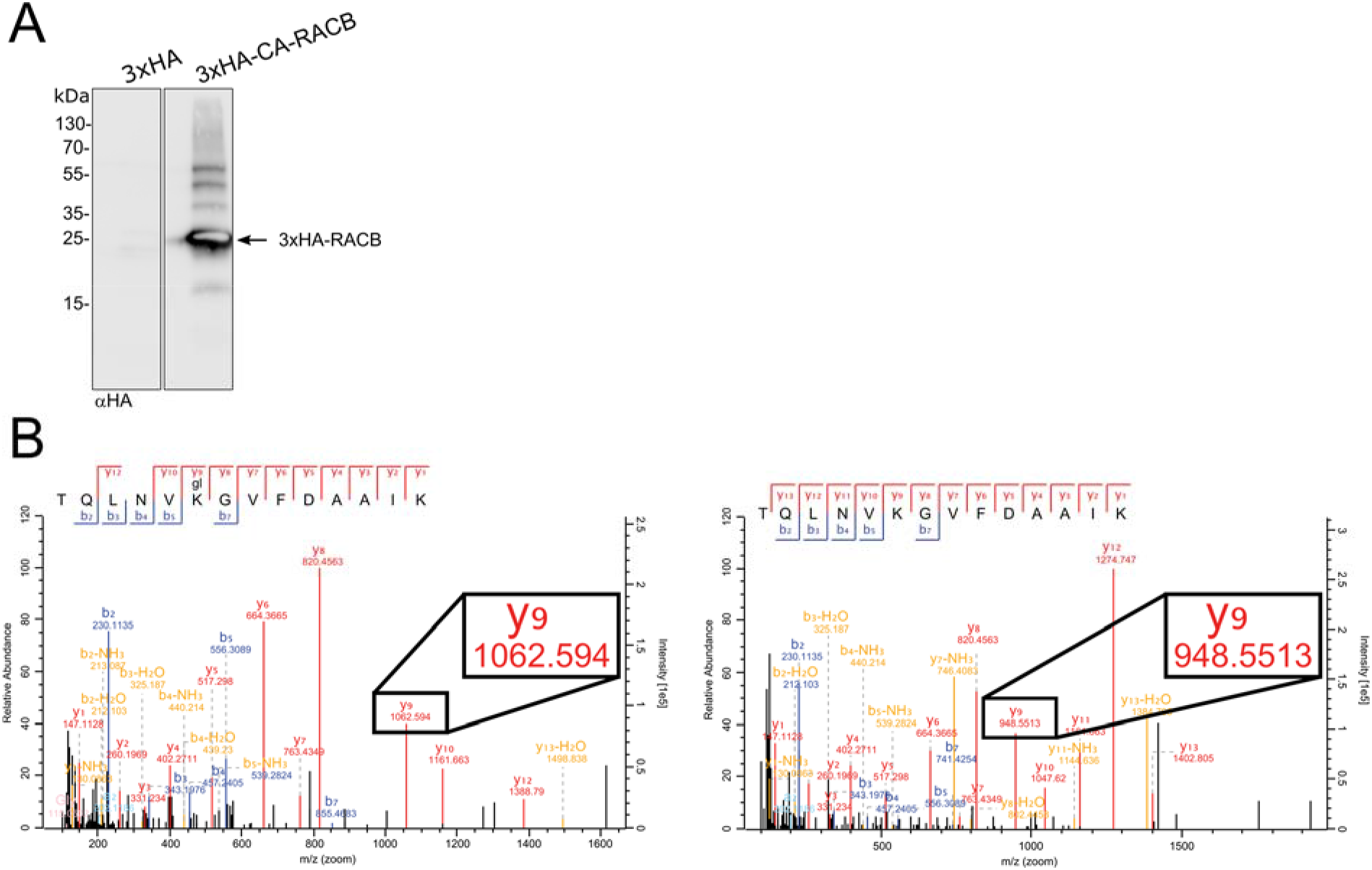
Constitutively activated RACB is ubiquitinated *in vivo*. **(A)** HA-tagged CA-RACB and higher molecular weight bands were detected in an αHA-Western blot after MG132 treatment and αHA-immunoprecipitation from whole barley leaves stably expressing 3xHA or 3xHA-CA-RACB (Fig S2B). Plant material was pooled from transgenic lines BG657 E3 and E4 for 3xHA-CA-RACB and BG659 E2 and E3 for 3xHA. **(B)** Ubiquitination of CA-RACB at K167 was identified by mass spectrometry in a workflow described in S2B Fig. The left graph shows spectra of the recovered ubiquitinated peptide, while the right graph shows the corresponding unmodified peptide. The black box illustrates one example of such a size difference: fragment “y9” was recovered with two different masses. The difference corresponds to that of 2 glycines, which is typical after a tryptic digest of ubiquitinated proteins (Peng et al., 2003). Example: y9_modified_ (1062.59 m/z) – y9_unmodified_ (948.55 m/z) = 114.04 m/z = 2 * glycine (57.02 m/z).

**Fig 6.**
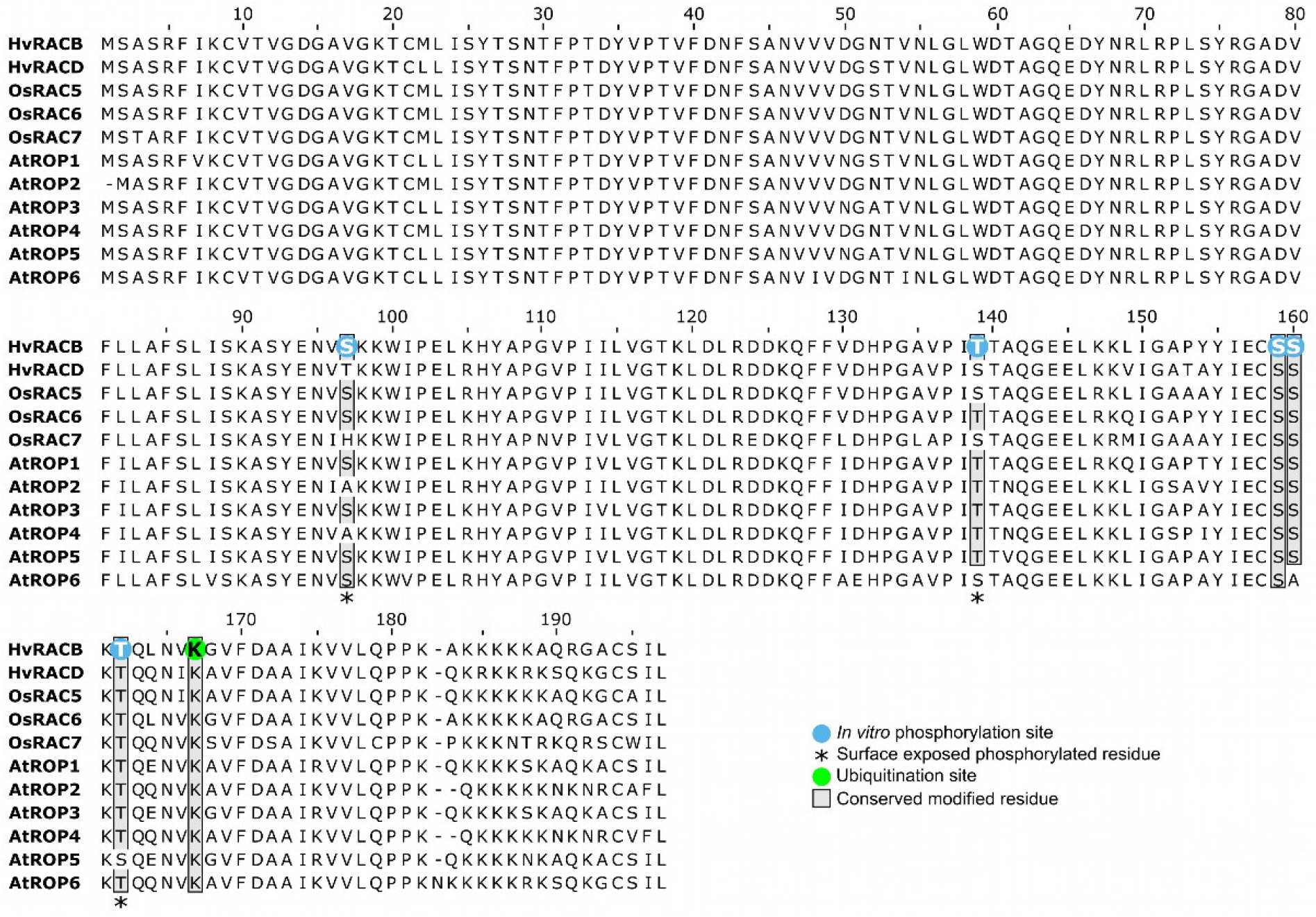
Alignment of type I ROPs from barley (*Hordeum vulgare*, *Hv*), rice (*Oryza sativa, Os*) and *Arabidopsis thaliana* (*At*). The ubiquitinated lysine residue in barley RACB (K167, green) is conserved across all type I ROPs from the three species (shaded box). *In vitro* phosphosites of RACB (blue) are conserved among some type I ROPs (shaded box). Surface-exposed phosphosites are highlighted with asterisks.

### RACB K167 regulates protein stability

Next, we wanted to know if RACB K167 plays a role in regulating protein stability. For this, we generated non-ubiquitinatable K167R mutants of the different RACB activity-variants (WT, CA, DN [dominant negative RACB-T20N (Schultheiss et al., 2003)]). We substituted lysine with arginine, because both amino acids are positively charged and structurally similar, but arginine is not recognized as an ubiquitin acceptor site. In order to assess if K167 influences protein stability in barley epidermal cells, we fused GFP to the N-terminus of RACB and compared the fluorescence levels of regular GFP-WT/CA/DN-RACB with those of their corresponding K167R mutants via confocal microscopy after biolistic transformation (Fig 7). Since fluorescence levels inherently vary between single transiently transformed cells, we co-expressed free mCherry as a normalizer in all experiments and used the calculated ratio of GFP to mCherry fluorescence for comparisons. Thereby, we measured that the K167R mutants of GFP-CA- and GFP-DN-RACB showed stronger fluorescence signals than their original K167 counterparts, while the GFP-WT-RACB(-K167R) fluorescence levels displayed no difference. Comparison of total protein abundance between RACB variants was not feasible, because laser excitation levels had to be adjusted for imaging of WT/CA/DN-RACB.

**Fig 7.**
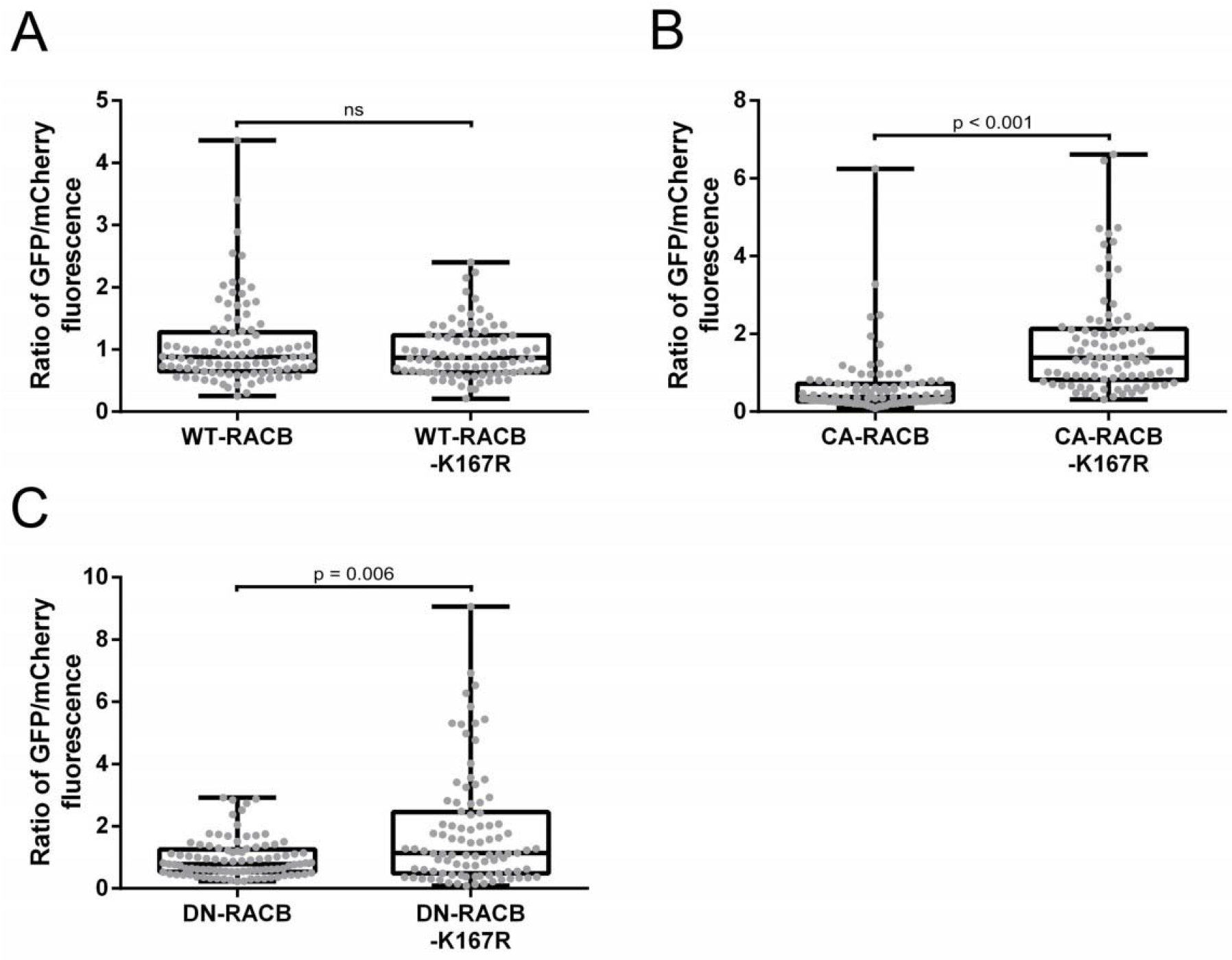
Quantification of GFP-RACB(-K167R) fluorescence intensities. GFP-tagged RACB-variants (WT **(A)**, CA **(B)** and DN **(C)**) and their corresponding non-ubiquitinatable K167R mutants were transiently expressed in single barley epidermal cells via particle bombardment. Free mCherry was co-expressed in all experiments as a normalizer. Mean pixel fluorescence intensities for GFP and mCherry were measured using a confocal laser-scanning microscope. For comparison, ratios of fluorescence were calculated by dividing the measured fluorescence signal of GFP-RACB by that of free mCherry in each individual cell. In case of GFP-CA- and GFP-DN-RACB, the K167R mutants displayed higher fluorescence levels than their non-mutated K167 counterparts, whereas GFP-WT-RACB(-K167R) fluorescence levels were similar. Each datapoint represents one transformed cell. All datapoints were collected over three biological replicates. Laser excitation levels were different for all three RACB activity variants, but kept identical for regular and K167R RACB forms. Statistical analysis was performed with Wilcoxon rank-sum tests with continuity correction against a p-value of 0.05.

To confirm that the RACB-K167R mutants are properly folded proteins and signaling competent, we tested if they are on one hand able to recruit downstream interactors to the plasma membrane and on the other hand influence the susceptibility of barley in favor of *Bgh*. We exclusively used CA-RACB in these experiments, since only this form of RACB was shown to govern signaling processes (Schultheiss et al., 2003; Pathuri et al., 2008; Schultheiss et al., 2008). In order to check if CA-RACB-K167R is promoting susceptibility to fungal cell entry in the barley-*Bgh*-interaction, we transiently overexpressed untagged CA-RACB(-K167R) or an empty vector control in barley epidermal cells before challenge inoculation with *Bgh* (Fig 8). At 40 hpi, we recorded a 39.4% or 34.6% increased susceptibility towards fungal penetration in cells overexpressing CA-RACB or CA-RACB-K167R, when compared to the control. Hence, we detected no difference in support of fungal penetration efficiency between CA-RACB and CA-RACB-K167R. For testing possible recruitment of a known RACB downstream interactor, we chose RIC171 because it was shown before to enrich at the cell periphery or plasma membrane when CA-RACB was co-expressed (Schultheiss et al., 2008). We used mCherry-RIC171 and GFP-CA-RACB(-K167R) fusion proteins for confocal microscopy in transiently transformed barley epidermal cells (Fig 9). Free CFP was co-expressed in all combinations as a marker for cytoplasmic and nuclear fluorescence. We observed that mCherry-RIC171 was strongly recruited towards the cell periphery in the presence of both GFP-CA-RACB and GFP-CA-RACB-K167R. No obvious recruitment of mCherry-RIC171 was detectable when free GFP was co-overexpressed and mCherry-RIC171 localized to the cytoplasm and nucleoplasm in this case.

**Fig 8.**
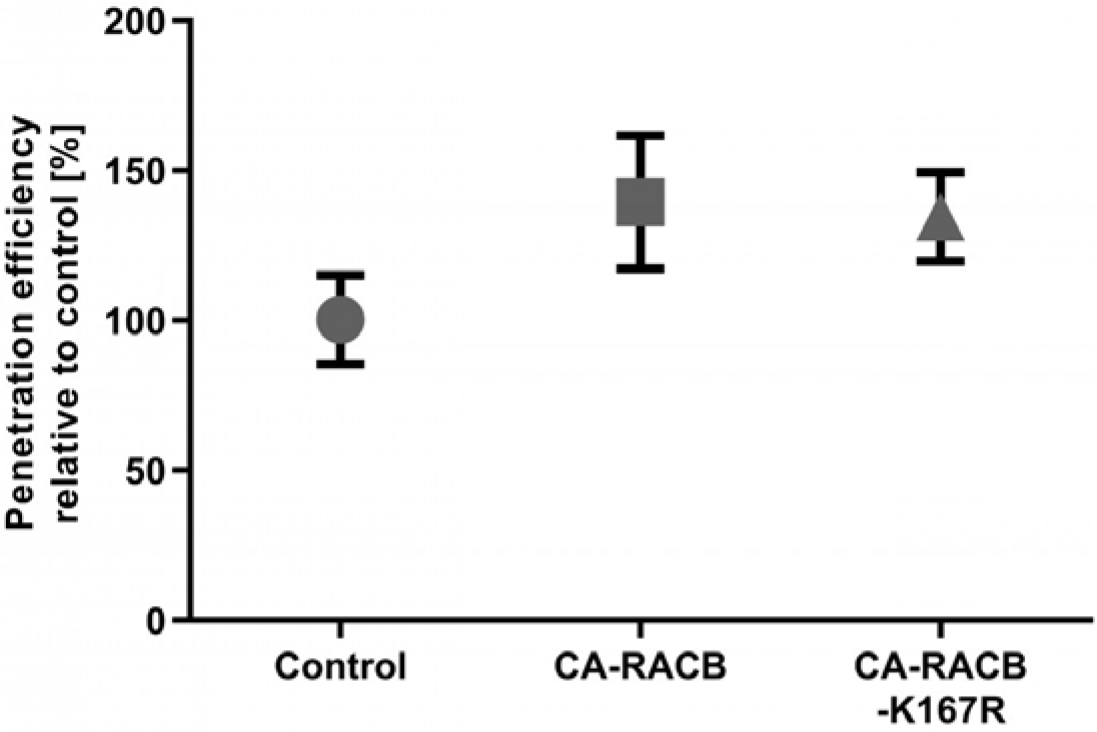
CA-RACB-K167R, similar to CA-RACB, increases barley susceptibility towards *Bgh*. Transient overexpression of untagged CA-RACB and CA-RACB-K167R in barley epidermal cells resulted in increased susceptibility towards *Bgh* when compared to the empty vector control. Transient transformation was achieved using particle bombardment. Fungal susceptibility was assessed 40 hpi. Mean *Bgh* penetration efficiency and standard error of the mean of 5 fully independent biological replicates are shown. Each datapoint represents one biological replicate, in which the penetration efficiency of *Bgh* was calculated using a minimum of 50 individual cells. Only single transformed barley epidermal cells were considered for analysis.

**Fig 9.**
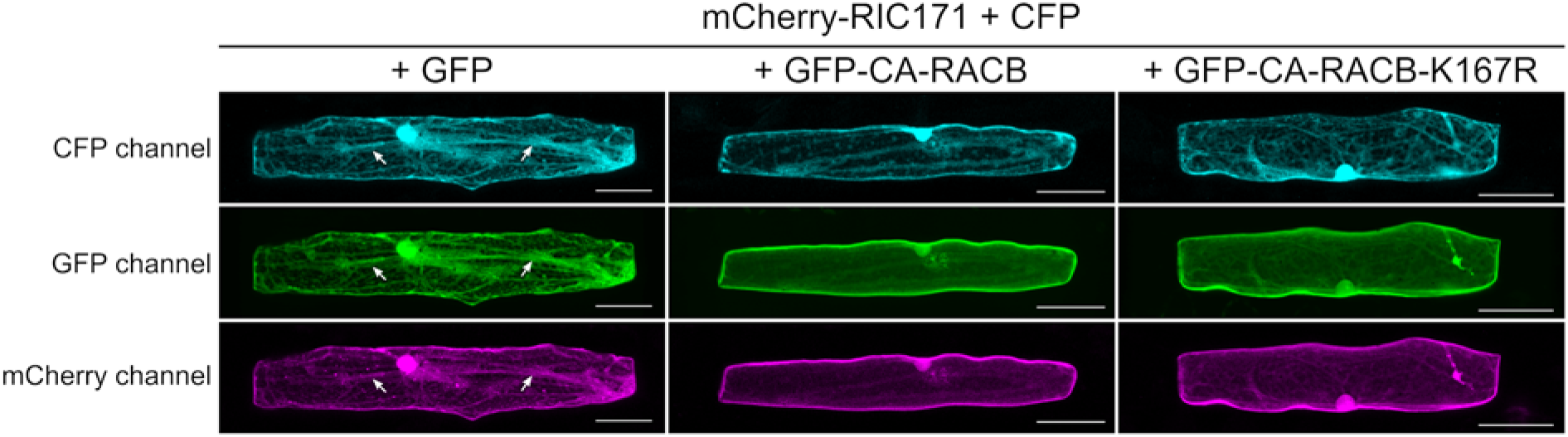
Recruitment of mCherry-RIC171 by GFP-CA-RACB-K167R. Localization of the mCherry-tagged RACB-interactor RIC171 (Schultheiss et al., 2008) was imaged in the presence of either free GFP, GFP-CA-RACB or GFP-CA-RACB-K167R. Free CFP was co-expressed in all cells as a marker for cytosolic and nuclear localization. All proteins were expressed transiently in barley epidermal cells via particle bombardment. In the absence of GFP-CA-RACB, mCherry-RIC171 fluorescence co-localized with GFP and CFP in the cytoplasm and nucleus (left panels, white arrows indicate cytosolic transvacuolar strands). When GFP-CA-RACB (center panels) or GFP-RACB-CA-K167R (right panels) were co-expressed, mCherry-RIC171 re-localized towards the periphery of the cell and showed only background fluorescence in the cytoplasm and nucleus. Representative images of three independent experiments are shown. In each experiment 15 cells per combination were imaged. Images display Z-stack maximum intensity projections of slices with 2 µm step increments. Brightness of the images was enhanced by 20% post-scanning for better visibility. The white scale bar in the bottom right corner of each panel represents 50 µm.

## Discussion

The function of RHO family proteins as signaling hubs and master regulators of cell polarity must be tightly controlled. Posttranslational modifications of RHO proteins, which enhance or suppress RHO protein activity, such as lipid modifications, phosphorylation and ubiquitination, have been well described in recent years (Abdrabou and Wang, 2018). Other PTMs including glycosylation and adenylylation have been predominantly demonstrated to be caused by virulence effectors from pathogenic organisms that target RHO family proteins of their hosts (Worby et al., 2009; Jank et al., 2013).

The regulation of ROP activity in plants by GEFs and GAPs has been well documented and occurs via their influence on the nucleotide binding state of the G-protein (Wu et al., 2000; Berken et al., 2005; Klahre and Kost, 2006; Hoefle et al., 2011; Akamatsu et al., 2013). However, for plants there is little data available on ROP activity regulation via posttranslational modifications beyond prenylation and *S*-acylation (Yalovsky, 2015). In this study, we show *in vivo* ubiquitination of a plant RHO protein, and our data also suggest a biological role for this PTM. Using transgenic barley plants overexpressing a CA-RACB variant, we identified higher molecular weight RACB derivatives on a Western blot, reminiscent of a typical ubiquitination pattern as described for e.g. the plant pattern-recognition receptor flagellin sensing 2 (FLS2) (Gohre et al., 2008; Lu et al., 2011). Mass spectrometry analysis evidenced RACB K167 as an ubiquitin acceptor site, which is fully conserved across all ROPs of barley, rice and Arabidopsis (Fig 6 and S4 Fig). The presence of this particular lysine even in mammalian Rac1 and Rac3 (Zhao et al., 2013; Dong et al., 2014) suggests a highly conserved ubiquitination motif across kingdoms. Conservation of two other ubiquitination sites from mammalian RHO proteins in RACB suggests that general regulatory mechanisms for small G-proteins could be conserved between plants and animals. Albeit we couldn’t evidence RACB K8 and K148 being ubiquitinated in our experimental setup, the corresponding lysines in RhoA and Rac1 are targeted for ubiquitination (S1 Fig), which influences actin cytoskeleton dynamics and associated cellular processes, such as cell motility, in mammalian cells (Wang et al., 2003; Ozdamar et al., 2005; Torrino et al., 2011; Oberoi et al., 2012; Castillo-Lluva et al., 2013). Nonetheless, it is not yet known whether and under which conditions a modification of these conserved sites might occur in ROPs. Regarding RACB, an association with protein degradation was supported by mutating specifically RACB-K167 to a non-ubiquitinatable arginine in various GFP-tagged RACB activity-variants (WT, CA, DN). CA-RACB-K167R was drastically more abundant than CA-RACB in barley epidermal cells, whereas this was less pronounced, but consistent for DN-RACB-K167R. Interestingly, the abundance of WT-RACB was not significantly influenced by the K167R mutation, which might be explained by the fact that wildtype RACB can still shuttle between GEF/GAP-mediated ROP activation and deactivation, whereas ubiquitination might be a slower process that predominantly takes place if RACB is arrested in a nucleotide bound form. Alternatively, efficient deubiquitination might be restricted to regulated WT-RACB. Generally, activated GTP-RACB could be ubiquitinated as an additional means of the plant cell to terminate active G-protein signaling besides inactivation via hydrolysis of GTP. This seems reasonable and could be conserved across kingdoms, as a similar mechanism has been shown for mammalian GTP-Rac1, which is ubiquitinated at the corresponding K166 to remove it from signaling in cell motility (Zhao et al., 2013). On the other hand, the purpose of ubiquitination of signaling incompetent DN-RACB could be a simple disposal mechanism.

The presence or absence of the protein stability-regulating lysine doesn’t appear to impinge on RACB-mediated downstream signaling. Both ubiquitinatable and non-ubiquitinatable CA-RACB could recruit the interaction partner RIC171 to the cell periphery, which was described before (Schultheiss et al., 2008). Additionally, overexpression of the higher abundant CA-RACB-K167R showed the same increase in barley susceptibility towards penetration by *Bgh* as CA-RACB (Schultheiss et al., 2003; Schultheiss et al., 2008). Hence, the sheer abundance of activated RACB might not be limiting for fungal penetration success in a situation in which CA-RACB is overexpressed. However, ubiquitination could have an impact on RACB’s ability to interact with other proteins. For instance, non-ubiquitinated K167 is presumably bound by PRONE-GEFs and CRIB-domain-containing proteins, since it is in both cases part of the respective binding interface (see S5 Fig, (Abdul-Manan et al., 1999; Thomas et al., 2007; Schaefer et al., 2011)). Ubiquitination of K167 could hence sterically block these interfaces and therefore hinder activation of RACB by PRONE-GEFs and interaction with downstream CRIB-containing executors. In both scenarios, RACB signaling would be effectively inhibited. In turn, it is likely that interaction of activated RACB with CRIB-domain-containing executors, such as RIC proteins (Schultheiss et al., 2008; Engelhardt et al., 2019), would mask the ubiquitination site. This could not only lead to increased protein stability, but also secure RACB’s signaling activity during crucial processes. In any case, RACB’s upstream or downstream interactors may have the potential to interfere with ubiquitination of RACB and hence proteasomal degradation, but this remains to be tested in the future.

This ubiquitin-dependent regulation of RACB activity could also involve RBK1 and SKP1-like, since RNAi-mediated knock-down of both proteins was shown to have a similar effect on RACB abundance (Reiner et al., 2016). We previously hypothesized that RACB is regulated similarly to mammalian Rac1. Phosphorylation of Rac1 S71 by AKT leads to SCF^FBXL19^-mediated ubiquitination at Rac1 K166 and subsequent proteasomal degradation, which blocks Rac1 from signaling (Kwon et al., 2000; Chang et al., 2011; Zhao et al., 2013). We speculated that activated RACB might be phosphorylated by RBK1 and then targeted for ubiquitination and degradation by an SCF-complex containing SKP1L (Huesmann et al., 2012; Reiner et al., 2016). While we did find ubiquitination at RACB K167 (which corresponds to Rac1 K166, S1 Fig), we did not identify RACB S74 as a phosphorylation site although the motif was covered in our MS data set. One potential reason for this might be that RBK1 belongs to RLCK class VI_A (Jurca et al., 2008; Huesmann et al., 2012; Reiner et al., 2015), while AKT belongs to the AGC kinases (Rademacher and Offringa, 2012). Interestingly, ROPs and AGCs also work together in polar cell development. For instance, the AGCVIII kinase D6PK binds to specific membrane lipids and cooperates with RACB-like type I *At*ROP2/*At*ROP6 to form root hairs (Stanislas et al., 2015). Thus far, phosphorylation of ROPs during interaction with AGC kinases has not been described, but it remains possible that AGC kinases phosphorylate ROPs, with S74 being one of the potential targets.

In Arabidopsis and barley, ROPs and RBK1 or related RLCK VI_A-class kinases have partially opposing biological functions. In previous studies, we have shown that gene silencing of barley *RBK1* provoked the opposite disease phenotype as *RACB* and supports RACB protein abundance (Huesmann et al., 2012; Reiner et al., 2016). In Arabidopsis, the ROP-activated RBK1-homolog RLCK VI_A3 is similarly involved in pathogen resistance rather than susceptibility (Reiner et al., 2015). Enders and co-workers could show that *rbk1* knock-out plants displayed hypersensitivity towards auxin, while *atrop4* and *atrop6* mutants were defective in auxin sensing. Furthermore, they suggested that *At*ROP4 and *At*ROP6 might be direct phosphorylation targets of *At*RBK1 (synonym: RLCK VI_A4). They hypothesized that *At*RBK1-mediated phosphorylation negatively influences the activity of both ROPs (Enders et al., 2017). However, this remains to be shown, since phosphorylation was investigated *in vitro* and no phosphosites were mapped in this study. Nevertheless, this suggests a perhaps general mechanism, in which ROPs are negatively regulated by phosphorylation through RLCK VI_A members in distinct signaling branches. However, we would like to stress that *in vivo* phosphorylation of RACB by RBK1 could neither be proved nor disproved in this study.

The *in vitro* phosphosites S97, T139 and T162, which we identified in RACB, are predicted to be surface-exposed and therefore could theoretically be targets for RBK1-dependent phosphorylation of RACB *in vivo* (Fig 2). This is less clear for S159 and S160. Nevertheless, one can speculate that these serine residues could become accessible *in vivo* after other posttranslational modifications of RACB, which cause a conformational shift. Alternatively, these sites become more exposed during interaction with RBK1. The positions of the phosphosites within RACB suggest that the modification of particularly S159, S160 and T162 could interfere with RACB lipidation and nucleotide cycling, since these residues are next to the putative *S-*acylation site C158 and the nucleotide binding pocket (Fig 2 and S1 Fig; (Sorek et al., 2010; Sorek et al., 2017)). Furthermore, *in vivo* phosphorylation of all five residues could affect RACB activation, as they are part of or in direct proximity of a conserved PRONE-GEF binding interface (see S5 Fig, (Sørmo et al., 2006; Thomas et al., 2007; Fricke and Berken, 2009)). However, this remains a speculation at this point.

All five in *vitro* phosphorylated residues in RACB are partially conserved in other type I and type II ROPs, indicating that these sites might be common targets for phosphorylation. The only evidenced *in vivo* ROP phosphosite, S138 from *At*ROP10 and *At*ROP11 (S4 Fig; (Mergner et al., 2020)), is not conserved in RACB, which indicates that this might be a type II ROP-specific modification site. In any case, introducing point mutations at these sites could help answering which role they play in regulating ROP function.

Comparison of RACB phosphosites with those of mammalian Rac1 and RhoA revealed that there is no overlap between mapped animal and plant phosphosites, although several modified residues are conserved across kingdoms (S1 Fig). Thus, it is well possible that RACB is also phosphorylated at Y67, S74 and S89 by other kinases. Still, conservation of these amino acids suggests that RACB could be regulated in a similar fashion as mammalian RHO family proteins. Interestingly, phosphorylation of these residues in mammalian RHO proteins directly and indirectly regulates actin cytoskeleton dynamics (Chang et al., 2011; Zhao et al., 2013; Tong et al., 2016), a process also influenced by RACB in barley (Opalski et al., 2005). Therefore, it remains conceivable that phosphorylation of these conserved sites may be important for ROP regulation during plant cytoskeleton rearrangement.

Using stable transgenic plants, our data further support that posttranslational prenylation may be important for the signaling ability of RACB. Full-length CA-RACB increased barley susceptibility towards *Bgh*, whereas the truncated CA-RACB-ΔCSIL mutant did not. We have shown before in transiently transformed barley cells that removal of the CaaX-box prenylation motif CSIL from RACB results in a shift of CA-RACB’s subcellular localization from the plasma membrane to the cytosol and abolishes function in susceptibility (Schultheiss et al., 2003). We have also reported that the interaction of RACB with its downstream effectors mostly takes place at the plant plasma membrane (Schultheiss et al., 2008; Hoefle et al., 2011; McCollum et al., 2020). Our data with CSIL truncations in stable transgenic barley provide now another piece of evidence that ROPs need to be membrane-localized to perform their function, as it has been described before for Arabidopsis (Sorek et al., 2011; Yalovsky, 2015).

## Conclusion

Regulation of RHO family proteins by PTMs has long been established in the animal research field. In plants, however, we are only at the beginning of discovering these regulatory mechanisms of PTMs in ROP signaling. Here, we present data on *in vivo* ubiquitination of a barley ROP. The identified ubiquitination site is conserved in all ROPs from barley, rice and Arabidopsis, and has been described before for mammalian RACB-homologs. This suggests that the lysine residue corresponding to RACB K167 is a general target for ubiquitination across kingdoms and that this regulates protein abundance.

## Experimental procedures

### Heterologous protein expression, purification, and kinase activity measurements

A cDNA clone of CA-*RACB* ((Schultheiss et al., 2003), NCBI GenBank identifier for wildtype RACB: CAC83043.2) was cloned into the bacterial expression vector pET26b out of frame of the N-terminal pelB, but in frame with a C-terminal 6xHIS (Novagen, Merck KGaA, Darmstadt, Germany). This way, no potential artificial phosphorylation site was included (Dorjgotov et al., 2009). The barley protein *RBK1* (NCBI GenBank identifier: HE611049.1) was cloned into the pET28a vector in frame with an N-terminal 6xHIS. Cloning was achieved by restriction-ligation after PCR amplification of the cDNA inserts using *Eco*RI and *Xho*I flanking restriction sites on the C- and N-terminal sequence-specific primers. CA-*RACB* was amplified using RACBCA-EcoRI-F + RACBCA-XhoI-R primers, while both wildtype and kinase-dead *RBK1* were amplified with RBK1-F + RBK1-R primers. The kinase dead (KD, K245D) mutant of *RBK1* was constructed by PCR-mediated site-directed mutagenesis using primers RBK1KD-mut-F + RBK1KD-mut-R. After PCR, the parental DNA was digested with *Dpn*I and the whole reaction was transformed into *E. coli* DH5α cells. Following transformation, plasmids were extracted using a routine Miniprep protocol and checked for mutation by sequencing. All primers can be found in S5 Table.

Proteins were purified using HisSelect Sepharose beads (Sigma-Aldrich, St. Louis, USA) and the *in vitro* kinase reactions were carried out as described earlier (Huesmann et al., 2012; Reiner et al., 2016). Two parallel reactions, one with and one without radioactive γ-^32^P-ATP were performed with both WT and KD RBK1, using the same amount of RACB-6xHIS as substrate. The amount of the proteins in the reactions was app. 150 pmol of RACB-6xHIS with 4 pmol of 6xHis-RBK1-WT or 12 pmol 6xHis-RBK1-KD. The radioactive reaction for autoradiography contained 10 µM of cold ATP and 0.2 MBq γ-^32^P-ATP, while 100 µM ATP were included in the non-radioactive reaction destined for mass spectrometry analysis. The reactions were terminated after 2.5 h and proteins were separated on 12% SDS-polyacrylamide gels that were stained by Coomassie Brilliant Blue using standard methods. Radioactive gels were dried and exposed to x-ray films. From the non-radioactive gel, the band containing RACB-6xHIS was cut out and sent for phosphosite mapping.

To show stability of both WT and KD 6xHis-RBK1, a routine Ponceau S stain was perfomed.

### Bioinformatical analysis

The putative RACB protein fold was built by homology-modelling using SWISS-MODEL (Waterhouse et al., 2018) with GDP-bound AtROP4 (PDB entry: 2NTY.1 (Thomas et al., 2007)) as template for GDP-bound RACB and GNP-associated *Hs*RAC1 as template for GTP-RACB (PDB entry: 3TH5.1 (Krauthammer et al., 2012)). SWISS-MODEL GMQE (global model quality estimate) scores were 0.75 for GDP-RACB and 0.66 for GTP-RACB, and QMEAN DisCo (qualitative model energy analysis with consensus-based distance constraint) (Studer et al., 2020)) scores were 082 ± 0.07 and 0.80 ± 0.07, respectively. The final RACB models contained amino acids 1-178 for GDP-RACB and 7-178 for GTP-RACB (from 197 total RACB amino acids). PyMOL V2.3.5 (Schrodinger, 2015) was used to generate protein surface structure and cartoon illustrations.

All protein sequence alignments were done in Jalview V2.11.0 (Waterhouse et al., 2009) with the MAFFT algorithm running at default settings (Katoh and Toh, 2010). NCBI GenBank identifiers for all aligned proteins are:

HvRACB: CAC83043.2; HvRACD: CAD27895.1; HvRAC1: CAD57743.1; HvRAC3: BAJ98596.1; HvROP4: CAD27896.1; HvROP6: CAD27894.1; OsRAC1: XP_015621645.1; OsRAC2: XP_015638759.1; OsRAC3: XP_015625155.1; OsRAC4: XP_015641323.1; OsRAC5: XP_015627011.1; OsRAC6: XP_015625732.1; OsRAC7: XP_015627590.1; AtROP1: NP_190698.1; AtROP2: NP_173437.1; AtROP3: NP_001077910.1; AtROP4: NP_177712.1; AtROP5: NP_195320.1; AtROP6: NP_001190916.1; AtROP9: NP_194624.1; AtROP10: NP_566897.1; AtROP11: NP_201093.1; HsRAC1: NP_001003274.1; HsRHOA: 5FR2_A.

### Generation of transgenic plants

A full-length cDNA clone of CA-*RACB* (carrying the G15V mutation as described in (Schultheiss et al., 2003)) and CA-*RACB*-ΔCSIL (lacking the last 4 amino acids, but with an artificial stop codon) were amplified by PCR using primers RACBCA-Esp-Start + RACBCA-Esp-Stop and RACBCA-Esp-Start + RACBCA-ΔCSIL-Esp-Stop, respectively. Both constructs were cloned first into pGGEntL-EP12 and subsequently into pGGInAE-226n_35SP_Ntag-3xHA_35ST vectors using procedures adapted from Golden Gate-based cloning (Engler et al., 2008), generating the 3x*HA*-CA-*RACB*(-ΔCSIL) fusion constructs. From these vectors, the CA-*RACB*(-ΔCSIL) fusion constructs were re-amplified by PCR using primers 3xHA-XmaI-F and RACBCA-XmaI-R (3x*HA*-CA-*RACB*) and 3xHA-XmaI-F + RACBCA-ΔCSIL-XmaI-R (3x*HA*-CA-*RACB*-ΔCSIL). The free 3x*HA*-control was amplified using primers 3xHA-XmaI-F + UL1-XmaI-R. All constructs were introduced into pUbifull-AB-M, a derivative of pNOS-AB-M (DNA-Cloning-Service, Hamburg, Germany) with the maize *POLYUBIQUITIN 1* promoter being inserted in the multiple cloning site, by restriction-ligation using *Xma*I. From there, *Sfi*I fragments were transferred into the generic binary vector p6i-2×35S-TE9 (DNA-Cloning-Service, Hamburg, Germany). The binary plasmids were transferred into *A. tumefaciens* strain AGL1 (Lazo et al., 1991) and transgenic plants were generated by agro-inoculation of immature barley embryos according to Hensel et al. (2009). Transgenic plants were identified by *HYGROMYCIN PHOSPHOTRANSFERASE*-specific PCR, which yielded 47 lines for 3xHA-CA-RACB (BG657 E1-E49; E7 and E41 were not transgenic), 37 lines for 3xHA-CA-RACB-ΔCSIL (BG658 E1-E39; E19 and E38 were not transgenic) and 40 lines for 3xHA (BG659 E1-E42; E17 and E37 were not transgenic). All lines were propagated under normal growth conditions in the greenhouse. Since all descending lines are segregating in generation T_1_, individual plants were screened for transgene integration. This was achieved using Western blotting detecting the HA-tag (antibody: Anti-HA-Peroxidase, Sigma-Aldrich, St. Louis, USA, 12013819001) and resistance against hygromycin-induced discoloration. For the latter, 7 d old leaves were detached and placed on 0.8% water-agar plates containing 200 µg/ml Hygromycin B (Roth, Karlsruhe, Germany, CP12.2). These plates were put into growth chambers operating under long-day conditions (see below) for roughly 4 d, until discoloration of hygromycin non-resistant plants was observed. After screening, ten positive plants of two lines per construct were picked for an additional round of propagation: we chose BG657 E3 and E4 for 3x*HA*-CA-*RACB*, BG658 E4 and E9 for 3x*HA*-CA-*RACB*-ΔCSIL and BG659 E2 and E3 for 3x*HA*. Since the offspring of these lines remained segregating in generation T_2_ and no homozygous plants were obtained, all descendants were screened for transgene expression as described above before being used in any experiment. All primers can be found in S5 Table.

### Plant and fungal growth conditions

Transgenic barley (*Hordeum vulgare*) plants of spring-type cultivar Golden Promise were grown under long-day conditions (16 h light, 8 h darkness) with a photon flux of at least 150 µmol m^−2^ s^−1^ at 18 °C and a relative humidity of 65%. Transgene-expressing plants of generation T_2_ were used in all experiments but one. An exception was made for the *Bgh* penetration assay (see below), because propagation of the T_2_ plants was not ready at that time. Therefore, we used corresponding T_1_ plants in this experiment. For all other experiments, pools of T_2_ plants expressing the same construct were used. For 3x*HA*-CA-*RACB*, we pooled plants of lines BG657 E3 and E4, BG658 E4 and E9 for 3x*HA*-CA-*RACB*-ΔCSIL and BG659 E2 and E3 for 3x*HA*. For transient transformation experiments (see below) and propagation of the barley powdery mildew fungus *Blumeria graminis* f.sp. *hordei* race A6 (*Bgh*), non-transformed wildtype barley cultivar Golden Promise plants were grown under the same growth conditions.

### Assessment of *Bgh* penetration efficiency

Primary leaves of transgenic plants were cut and placed on 0.8% water-agar plates. These plates were inoculated with 100-130 spores/mm^2^ of *Bgh* and placed under normal growth conditions for 40 h, followed by fixing and destaining leaves in 70% (v/v) ethanol. Fungal spores and appressoria were stained with an ink solution (10% (v/v) Pelikan Ink 4001® (Pelikan, Hannover, Germany), 25% (v/v) acetic acid) and analyzed by transmission light microscopy. Haustorium establishment was counted as successful penetration, while stopped fungal growth after appressorium formation was counted as successful plant defense. For each construct, three transgene-expressing plants of one T_1_ line (BG657 E4 for 3x*HA*-CA-*RACB*, BG658 E4 for 3x*HA*-CA-*RACB*-ΔCSIL and BG659 E2 for 3x*HA*) were analyzed, with at least 96 evaluated plant-fungus interaction sites per plant.

For the analysis of *Bgh* susceptibility after transient overexpression of untagged CA-RACB(-K167R), a full-length cDNA clone of *RACB* was first amplified with Gateway® attachment sites using primers RACB-Gateway-F + RACB-Gateway-R and cloned into the Gateway® entry vector pDONR223 using routine Gateway® BP-cloning procedures (Invitrogen, Carlsbad, California, USA). From pDONR223, *RACB* was transferred into the Gateway®-compatible plant expression vector pGY1-GW (generated in Engelhardt et al. (2019)) using routine Gateway® LR-cloning. A CA-*RACB*-K167R clone was generated from pGY1-*RACB* using subsequent site-directed mutagenesis with primers RACB-SDM-G15V-F + RACB-SDM-G15V-R and RACB-SDM-K167R-F + RACB-SDM-K167R-R. Integrity of the sequences and mutations was confirmed via Sanger sequencing. All primers can be found in S5 table.

For transient transformation via particle bombardment, three primary leaves of 7 d old wildtype barley plants (cultivar Golden Promise) were placed on 0.8% water-agar plates and transformed using the PDS-1000/He™ system (Bio-Rad, Hercules, California, USA). For one transformation reaction, 11 µl of 25 mg/ml spherical gold particles (diameter 1 µm, Nanopartz, Loveland, USA) were mixed with 1 µg plasmid DNA coding for pGY1-CA-*RACB*, pGY1-CA-*RACB*-K167R or pGY1 (empty) and 0.5 µg plasmid DNA coding for the transformation marker pUbi-*GUSplus* (a gift from Claudia Vickers, Addgene plasmid #64402, http://n2t.net/addgene:64402, (Vickers et al., 2003)). Afterwards, 15 µl CaCl_2_ (to a final concentration of 0.5 M) and 7.5 µl of 20 mg/ml Protamine (Sigma-Aldrich, St. Louis, USA) were added and the solution was incubated at RT for 30 min, with occasional mixing. Subsequently, the solution was washed twice, first with 500 µl 70% ethanol and second with 500 µl 100% ethanol. Finally, the gold particles were dissolved in 6 µl 100% ethanol using a sonicator and used in the PDS-1000/He™ System. For each plasmid combination, nine barley leaves distributed on three plates were transiently transformed per biological replicate. After transformation, all plates were placed in a growth chamber operating under the above stated conditions. These plates were then challenge-inoculated with 100-130 spores/mm^2^ of *Bgh* at 24 h post transformation and incubated under normal plant growth condition for 40 h. Afterwards, the leaves were first transferred into plates containing a GUS-staining solution (0.1 M Na_2_HPO_4_/NaH_2_PO_4_ buffer pH 7, 0.01 M sodium EDTA, 0.005 M potassium hexacyanoferrat (II), 0.005 M potassium hexacyanoferrat (III), 0.1% (v/v) Triton X-100, 20% (v/v) methanol, 0.5 mg/ml X-gluc (1,5-bromo-4-chloro-3-indoxyl-β-D-glucuronic acid, cyclohexylammonium salt (Carbosynth, Bratislava, Slovak Republic), (Schweizer et al., 1999)), vacuum infiltrated for 3 min and incubated at 37 °C for 6 h in the dark, before finally being destained in 70% ethanol. For analysis of fungal penetration efficiency, fungal material was stained using a calcofluor staining solution (0.3% (w/v) Calcufluor-white, 50 mM Tris pH 9) before being subjected to light microscopy. Briefly, leaves were screened for transformed cells, which could be distinguished due to their blue GUS-stain. If a fungal attack occurred in a transformed cell, established haustoria were counted as successful penetration, while halted fungal growth after appressorium formation was counted as successful plant defense. The *Bgh* penetration efficiency for each plasmid combination was calculated by dividing the number of successful penetration events by the sum of all successful penetration and defense events in this combination. Five biological replicates of this experiment were analyzed, with at least 50 evaluated plant-fungus interactions per plasmid combination per replicate. Fig 8 displays all *Bgh* penetration efficiencies divided by the mean penetration efficiency of all empty vector controls with the respective standard error of the mean. All penetration efficiency data (stable and transient assays) were visualized with GraphPad Prism version 6. Statistical analysis was performed in RStudio V1.2.5033 (source: https://rstudio.com/) using R V3.6.3 (source: https://www.R-project.org). Normal distribution and homogeneity of variance in the datasets were confirmed with Shapiro-Wilk’s test and Levene’s test, respectively. Significant differences of results were assessed using a One-way ANOVA with Tukey’s HSD against a p-value of 0.05.

### Protoplast transformation, proteasome inhibition and protein extraction

Transgenic barley protoplasts stably expressing 3xHA-CA-RACB (pools of lines BG657 E3 and E4) or 3xHA (pools of lines BG659 E2 and E3) were isolated and super-transformed as described in Reiner et al. (2016), with the following deviations: transformation reactions were scaled to 1 ml of protoplasts and overnight incubation was conducted in 1 ml buffer WI in single cell culture tubes. For transformation, 50 µg/ reaction of a plasmid coding for *GFP-RBK1* (from Huesmann et al. (2012)) or water (as a mock control) were used. Ten reactions per combination (S2A and S2B Figs) were prepared. After overnight incubation, MG132 was added to a final concentration of 10 µM, followed by incubation in the dark for 3 h. Protoplasts of corresponding reactions were pooled and pelleted by centrifugation at 120 g. Supernatants were removed and protoplasts were lysed by brief vortexing and snap-freezing in liquid nitrogen. Proteins were resuspended in a modified extraction buffer (10% (w/v) glycerol, 25 mM Tris pH 7.5, 150 mM NaCl, 10 mM DTT, 1 mM PMSF, 1% sodium deoxycholate, 1x protease inhibitor (Sigma-Aldrich, St. Louis, USA, P9599), 1 tablet /10 ml PhosSTOP™ (Sigma-Aldrich, St. Louis, USA, 4906845001)) and incubated on ice for 30 min with occasional mixing. This solution was centrifuged at 4 °C to pellet cell debris, the supernatant was digested with trypsin and used in phosphopeptide enrichment (S2A Fig (center panel) and S3B Fig). In one experiment, where the phosphopeptide enrichment was skipped (S2A Fig (bottom panel)), proteins were subjected to immunoprecipitation (see below), loaded on a SDS-PAGE and then submitted to in-gel tryptic digest prior to mass spectrometry. All buffers described in the immunoprecipitation paragraph were used. In another experiment (S2A Fig (top panel)), protoplast lysates were subjected to immunoprecipitation using a modified extraction buffer (10% (w/v) glycerol, 25 mM Tris pH 7.5, 150 mM NaCl, 10 mM DTT, 1 mM PMSF, 50 μM MG132, 1x protease inhibitor (Sigma-Aldrich, St. Louis, USA, P9599), 1 tablet /10 ml PhosSTOP™ (Sigma-Aldrich, St. Louis, USA, 4906845001)) and washing buffer (10% (w/v) glycerol, 25 mM Tris pH 7.5, 150 mM NaCl, 1 mM PMSF, 50 μM MG132, 1x protease inhibitor (Sigma-Aldrich, St. Louis, USA, P9599), 1 tablet /10 ml PhosSTOP™ (Sigma-Aldrich, St. Louis, USA, 4906845001)) and used for on-bead tryptic digest prior to phosphopeptide enrichment via IMAC.

### Immunoprecipitation and Western blotting

In order to show protein stability of 3x*HA*-CA-*RACB* and 3x*HA*-CA-*RACB*-ΔCSIL, we performed a routine αHA-immunoprecipitation from untreated transgenic leaves (lines BG657 E3 and E4 and BG658 E4 and E9, respectively), followed by an anti-HA Western blot (see below). For ubiquitination analysis, transgenic barley leaves expressing 3xHA-CA-RACB (lines BG657 E3 and E4) or 3xHA (lines BG659 E2 and E3) were vacuum infiltrated for 3 min at -26 mm in Hg with a leaf-floating solution (0.45 M mannitol, 10 mM MES pH 5.7, 10 mM CaCl_2_, 0.0316 g /10 ml Gamborg B5 medium including vitamins (Duchefa, RV Haarlem, The Netherlands, G0210), 50 µM MG132). To determine, whether the observed ubiquitin-like laddering pattern of 3xHA-CA-RACB is MG132-dependent, we floated transgenic barley leaves expressing 3xHA-CA-RACB (lines BG657 E3 and E4) either on the float solution described above or in a solution that contained the equal volume of DMSO instead of MG132. After incubation in the dark for 3 h, leaves were snap-frozen in liquid nitrogen and homogenized using a TissueLyser II (QIAGEN, Hilden, Germany). Proteins were resuspended in extraction buffer (10% (w/v) glycerol, 25 mM Tris pH 7.5, 1 mM EDTA, 150 mM NaCl, 10 mM DTT, 1 mM PMSF, 0.5% Nonidet P40 substitute, 1x protease inhibitor (Sigma-Aldrich, St. Louis, USA, P9599)) by tumbling end-over-end at 4 °C for 30 min. Cell debris was removed by centrifugation and the supernatants were used for immunoprecipitation (IP). IPs were conducted according to routine procedures. Briefly, for each sample, 50 µM Pierce™ Anti-HA Magnetic Beads (Thermo Fisher Scientific, Waltham, USA) were equilibrated 3x in extraction buffer before adding supernatants. HA-tagged proteins were immunoprecipitated by tumbling end-over-end for 1 h at 4 °C, then washed 5x with washing buffer (10% (w/v) glycerol, 25 mM Tris pH 7.5, 1 mM EDTA, 150 mM NaCl, 1 mM PMSF, 1x protease inhibitor (Sigma-Aldrich, St. Louis, USA, P9599)). For selected RACB phosphopeptide screenings (S2A Fig), the beads were directly prepared for mass spectrometry. Otherwise, proteins were eluted in 100 µl of 2x NuPAGE™ LDS Sample Buffer (Thermo Fisher Scientific, Waltham, USA, NP0008) and boiled at 95 °C for 10 min. After the IP, 25 µl of eluate were separated on a 12% SDS-PAGE and proteins were transferred to a PVDF membrane. HA-tagged proteins were detected with an Anti-HA-Peroxidase antibody (Sigma-Aldrich, St. Louis, USA, 12013819001). For the RACB ubiquitination screening, the remaining 75 µl were loaded on another 12% SDS-PAGE and silver stained after separation. For this, the SDS-PAGE was incubated in a fixation solution (50% (v/v) methanol, 5% (v/v) acetic acid, 45% (v/v) water) for 1 h and subsequently washed with water three times (two times 2 min each, last time 1 h) on a shaking incubator. The gel was sensitized with a 0.02% (w/v) sodium thiosulfate solution for 2 min, then briefly washed with water twice. Afterwards, the gel was incubated at 4 °C in pre-chilled 0.1% (w/v) silver nitrate for 30 min, then rinsed two times with water for 30 s each. In order to stain proteins, the gel was incubated in a developing solution (0.04% (v/v) formaldehyde in 2% (w/v) sodium carbonate) until a sufficient degree of staining had been obtained. Then, the gel was washed five times in 1% (v/v) acetic acid to stop the reaction. Bands corresponding to 3xHA-CA-RACB and its higher molecular weight derivatives were excised from the gel and the gel slices were prepared for mass spectrometry. In order to show protein stability of 3x*HA*-CA-*RACB* and 3x*HA*-CA-*RACB*-ΔCSIL, we performed a routine HA-immunoprecipitation from untreated transgenic leaves (lines BG657 E3 and E4 and BG658 E4 and E9, respectively), followed by an αHA Western blot.

### Protein digest and phosphopeptide enrichment

In-gel digestion of SDS-PAGE gel segments was performed according to standard procedures. Briefly, excised gel segments were reduced and alkylated with 10 mM dithiothreitol (DTT) and 55 mM chloroacetamide (CAA) followed by digestion with trypsin (Roche). Dried peptides were reconstituted with 0.1% FA in water prior to LC-MS analysis. For peptide retention time monitoring, PROCAL peptide standard (Zolg et al., 2017) was added to peptide mixtures at a concentration of 100 fmol on column.

On-bead digestion of one immunoprecipitation sample (from the experiment described in S2A Fig (bottom panel)) was performed to facilitate a subsequent phosphopeptide enrichment. Dry beads-bound proteins were incubated with urea buffer (8 M Urea, 50 mM Tris HCl pH 8.5, 10 mM DTT, cOmplete^TM^ EDTA-free protease inhibitor cocktail (PIC) (Roche, Basel, Switzerland, 04693132001), Phosphatase inhibitor (PI-III; in-house, composition resembling Phosphatase inhibitor cocktail 1 (P2850), 2 (P5726) and 3 (P0044) from Sigma-Aldrich, St. Louis, USA) for one hour at 30 °C. The sample was alkylated (55 mM CAA), diluted 1:8 with digestion buffer (50 mM Tris-HCl pH 8.5, 1 mM CaCl_2_) and pre-digested with trypsin (1:100 protease:protein ratio) for 4 hours followed by overnight digestion with trypsin (1:100 protease:protein ratio) at 37 °C. The sample was acidified to pH 3 with TFA and centrifuged at 14,000 *g* for 15 min at 4°C. The supernatant was desalted on a 50 mg SepPAC column (Waters, Milford, USA; elution buffer 50% acetonitrile, 0.07% TFA).

Proteins from protoplast lysates were precipitated overnight with 10% trichloroacetic acid in ice-cold acetone at -20°C. Precipitates were washed two times with ice-cold acetone and dry samples re-suspended in urea buffer (8 M Urea, 50 mM Tris HCl pH 8.5, 1 mM DTT, cOmpleteTM EDTA-free protease inhibitor cocktail (PIC) (Roche, Basel, Switzerland, 04693132001), Phosphatase inhibitor (PI-III; in-house, composition resembling Phosphatase inhibitor cocktail 1 (P2850), 2 (P5726) and 3 (P0044) from Sigma-Aldrich, St. Louis, USA).

Protein concentration was determined with the Bradford assay (Bradford, 1976). 200 µg (experiment described in S2A Fig (center panel)) or 900 µg (all experiments described in S2B Fig) total protein was reduced (10 mM DTT), alkylated (55 mM CAA), diluted 1:8 with digestion buffer and pre-digested with trypsin (1:100 protease:protein ratio) for 4 hours followed by overnight digestion with trypsin (1:100 protease:protein ratio) at 37 °C. Samples were acidified to pH 3 with TFA and centrifuged at 14,000 *g* for 15 min at 4 °C. Supernatants were desalted on 50 mg SepPAC columns (Waters; elution buffer 50% acetonitrile, 0.07% TFA).

Phosphopeptide enrichment was performed using a Fe^3+^-IMAC column (Propac IMAC-10 4×50 mm, Thermo Fisher Scientific, Waltham, USA) as described previously (Ruprecht et al., 2017b). Shortly, desalted peptide samples were adjusted to 30% acetonitrile, 0.07% TFA. For quality control, 1.5 nmol of a synthetic library of phosphopeptides and their corresponding non-phosphorylated counterpart sequence (F1) (Marx et al., 2013), were spiked into samples prior to loading onto the column. The enrichment was performed with buffer A (0.07% TFA, 30% acetonitrile) as wash buffer and Buffer B (0.315% NH_4_OH) as elution buffer. Collected full proteome and phosphopeptide fractions were vacuum-dried and stored at -80°C until further use.

Phosphopeptides from the experiments described in S2A Fig (top and center panels) were re-suspended in 50 mM citrate, 0.1% formic acid (FA) and desalted on self-packed StageTips (five disks, Ø 1.5 mm C18 material, 3M Empore^TM^, elution solvent 0.1% FA in 50% ACN). Phosphopeptides from the experiment described in S2B Fig were re-suspended in 50 mM citrate, 0.1% formic acid (FA), fractionated on basic reverse phase StageTip microcolumns (five disks, Ø 1.5 mm C18 material, 3M Empore^TM^) and pooled to 4 fractions, as described previously (Ruprecht et al., 2017a). Dried peptides were reconstituted with 0.1% FA in water prior to LC-MS analysis.

*In vitro* kinase assay samples were mixed 1:4 with urea buffer (8 M urea, 50 mM Tris-HCl pH 7.5). Samples were reduced (5 mM DTT, 1h, room temperature), alkylated (15 mM CAA, 30 min, room temperature) and subsequently diluted 1:8 with digestion buffer. In-solution pre-digestion with trypsin (Roche, Basel, Switzerland) was performed for 4 hours at 37 °C (1:50 protease:protein ratio), followed by overnight digestion with trypsin (1:50 protease:protein ratio). Samples were acidified to pH 3 using trifluoroacetic acid (TFA) and centrifuged at 14,000 *g* for 5 min. The supernatant was desalted on self-packed StageTips (four disks, Ø 1.5 mm C18 material, 3M Empore^TM^, elution solvent 0.1% FA in 50% ACN). Dried peptides were reconstituted with 0.1% FA in water prior to LC-MS analysis.

### Mass spectrometry

Liquid chromatography-coupled mass spectrometry (LC-MS/MS) analysis was performed on Orbitrap mass spectrometer systems (Thermo Fisher Scientific, Waltham, USA) coupled on-line to a Dionex 3000 HPLC (Thermo Fisher Scientific, Waltham, USA). The liquid chromatography setup consisted of a 75 μm × 2 cm trap column packed in-house with Reprosil Pur ODS-3 5 μm particles (Dr. Maisch GmbH) and a 75 μm × 40 cm analytical column, packed with 3 μm particles of C18 Reprosil Gold 120 (Dr. Maisch GmbH, Ammerbuch, Germany). Peptides were loaded onto the trap column using 0.1% FA in water. Separation of the peptides was performed by using a linear gradient from 4% to 32% ACN with 5% DMSO, 0.1% formic acid in water over 60 min followed by a washing step (70 min total method length) at a flow rate of 300 nl/min and a column temperature of 50 °C. For phosphopeptide fractions, the gradient was adjusted to a two-step gradient from 4% to 27% ACN. The instruments were operated in data-dependent mode, automatically switching between MS and MS2 scans.

For the RACB ubiquitination screen samples (S2C Fig), full-scan mass spectra (m/z 360-1300) were acquired in profile mode on a Q Exactive HF with 60,000 resolution, an automatic gain control (AGC) target value of 3e6 and 10 ms maximum injection time. For the top 20 precursor ions, high resolution MS2 scans were performed using HCD fragmentation with 25% normalized collision energy, 30,000 resolution, an AGC target value of 2e5, 50 ms maximum injection time and 1.7 *m/z* isolation width in centroid mode. The underfill ratio was set to 1% with a dynamic exclusion of 20 s. To confirm and more precisely monitor the GlyGly- and phospho-modified peptides identified in the global analysis of these samples, a targeted parallel reaction monitoring assay (PRM) was set up on the Q Exactive HF using the same LC gradient settings. Peptide precursor charge states were chosen based on the previous identification results. In addition, a selection of PROCAL retention time calibration mixture peptides was monitored. PRM measurements were performed with the acquisition method switching between experiments after one duty-cycle. The first experiment consisted of a full scan MS1 spectrum recorded in the Orbitrap (360 to 1300 m/z, 15k resolution, AGC target 3e6, Maximum IT 50 ms). The second experiment consisted of a tMS2 PRM scan triggering MS2 scans based on a list containing retention time window, m/z and charge information. For the tMS2 PRM scan, the scheduled precursors were isolated (isolation window 1.3 m/z), fragmented via HCD (NCE 25%) and recorded in the Orbitrap (120 to 2,000 m/z, 15k resolution, AGC target 2e5, Maximum IT 100 ms).

For the kinase assay samples, full scan maximum injection time was 50 ms and MS2 scans were performed with a resolution of 15,000 and 100 ms maximum injection time. His-tagged CA-RACB and RBK1 peptides were set on an inclusion list and the dynamic exclusion set to 5s.

For the experiments described in S2B Fig, full-scan mass spectra (m/z 360-1300) were acquired in profile mode on a Q Exactive HF-X with 60,000 resolution, an automatic gain control (AGC) target value of 3e6 and 45 ms maximum injection time. For the top 20 precursor ions, high resolution MS2 scans were performed using HCD fragmentation with 26 % normalized collision energy, 30,000 resolution, an AGC target value of 2e5, and 1.3 *m/z* isolation width in centroid mode. The intensity threshold was set to 2.2e4 with a dynamic exclusion of 25 s. RACB and RBK1 peptides were set on an inclusion list. Full proteome samples were measured with 50 ms maximum injection time for MS2 scans, phosphopeptide samples with 100 ms.

For the experiment described in S2A Fig, full-scan mass spectra (m/z 360-1,300) were acquired in profile mode on an Orbitrap Fusion LUMOS with 60,000 resolution, an automatic gain control (AGC) target value of 4e5 and 50 ms maximum injection time. For the top 15 precursor ions, high resolution MS2 scans were performed using HCD fragmentation with 30% normalized collision energy, 30,000 resolution, an AGC target value of 5e4, 120 ms maximum injection time and 1.7 *m/z* isolation width in centroid mode. The intensity threshold was set to 2.5e4 with a dynamic exclusion of either 27 s (all peptides) or 10 s (RACB peptides set on an inclusion list).

Peptide and protein identification and quantification was performed with MaxQuant (Cox and Mann, 2008) using standard settings (version 1.5.8.3). Raw files were searched against a barley database (160517_Hv_IBSC_PGSB_r1_proteins_HighConf_REPR_annotation.fasta; IPK Gatersleben) and common contaminants. For the experiments described in S2B Fig, we added the phosphopeptide library sequences (Marx et al., 2013). For the kinase assay dataset, we used the recombinant protein sequences and an *E. coli* reference database (uniprot-proteome_UP000000625). Carbamidomethylated cysteine was set as fixed modification and oxidation of methionine, and N-terminal protein acetylation as variable modifications. Lysine ubiquitylation, phosphorylation of serine, threonine or tyrosine were set as variable modification were applicable. An m/z value of 115.0502 was specified as diagnostic ion for lysine glyglycylation (Zolg et al., 2018). Trypsin/P was specified as the proteolytic enzyme, with up to two missed cleavage sites allowed. The match between run function was enabled for matching samples and between fractions. Results were filtered to 1% PSM, protein and Site FDR. MS spectra were displayed with MaxQuant viewer.

RAW files from the PRM measurement of the ubiquitination screen samples were imported into Skyline (64-bit, v.4.1.0.11796) (MacLean et al., 2010) for data filtering and analysis and transition selection was manually refined to include site determining ions for each modification site. Peaks were integrated using the automatic peak finding function followed by manual curation of all peak boundaries and transitions. The summed area under the fragment ion traces data for every transition was exported for data visualization in Microsoft Excel and used to confirm the expected relative quantity of modified RACB peptides based on the gel segment. The mass spectrometry proteomics data have been deposited to the ProteomeXchange Consortium via the PRIDE (Vizcaíno et al., 2016) partner repository with the dataset identifier PXD019273.

### Confocal microscopy

For the generation of GFP-tagged RACB-variants, *RACB* coding sequence was mutated in pDONR223-*RACB* (from above) to create the G15V (CA), T20N (DN), K167R, G15V-K167R and T20N-K167R mutants using site-directed mutagenesis with the respective primer pairs RACB-SDM-G15V-F + RACB-SDM-G15V-R, RACB-SDM-T20N-F + RACB-SDM-T20N-R and RACB-SDM-K167R-F + RACB-SDM-K167R-R. Subsequently, the *RACB* mutants were transferred from pDONR223 into the Gateway®-compatible plant expression vector pGY1-*GFP* (described in Engelhardt et al. (2019)) via Gateway® LR-cloning. Integrity of the sequences and mutations were confirmed via Sanger sequencing. All primers can be found in S5 Table. Protein stability of GFP-tagged RACB and its variants (WT, CA, DN and WT-K167R, CA-K167R, DN-K167R) was indirectly assessed in transiently transformed barley epidermal cells using confocal laser-scanning microscopy. For this, three primary leaves of 7 d old wildtype barley plants (cultivar Golden Promise) were placed on 0.8% water-agar plates and transformed using biolistic bombardment with the PDS-1000/He™ system (Bio-Rad, Hercules, California, USA) as described above. For one transformation reaction, 1 µg of plasmid DNA coding for a GFP-tagged RACB variant and 0.5 µg of plasmid DNA coding for free mCherry were mixed. At 24 h post transformation, the leaves were analyzed using a Leica TCS SP5 CLSM mounted on a DM6000 stage (Leica, Wetzlar, Germany). Single transformed barley epidermal cells were identified and fluorophores were sequentially scanned. Hereby, GFP was excited with a 488 nm Argon laser and detected between 500 nm and 550 nm using a HyD detector, while mCherry was excited with a 561 nm laser line and detected between 570 nm and 620 nm using another HyD detector. To analyze and compare fluorescence intensities between one RACB variant and its K167R counterpart, appropriate laser and detector settings were chosen initially and kept constant for all cells and biological replicates of this combination. For all cells, Z-stacks with 2 µm step size were acquired, ranging from the top of the cell until the full nucleus was captured. The number of Z-stacks was kept constant for each RACB variant combination. For one biological replicate, at least 25 cells were imaged per construct, with three biological replicates in total. Single-cell mean pixel fluorescence intensities were measured separately for GFP and mCherry, as 8 bit grey scale values using Fiji (Schindelin et al., 2012), with regions of interest (ROIs) over one whole cell that was displayed a maximum intensity projection of the Z-stacks. Data points of three independent experiments were collected and graphed in GraphPad Prism version 6. Statistical analysis was performed in RStudio V1.2.5033 (source: https://rstudio.com/) using R V3.6.3 (source: https://www.R-project.org). Normal distribution and homogeneity of variance were tested with Shapiro-Wilk’s test and *Fisher’s*-test, respectively. Significant differences of results were assessed using Wilcoxon’s rank-sum test with continuity correction against a p-value of 0.05.

For the RIC171 recruitment assay, an mCherry-RIC171 clone was generated by first amplifying a coding sequence of *RIC171* (Schultheiss et al., 2008) via PCR with the primer pair RIC171-Gateway-F + RIC171-Gateway-R and cloning it into the Gateway® entry vector pDONR223 using Gateway® BP-cloning. From there, RIC171 was transferred into the Gateway®-compatible plant expression vector pGY1-nmCherry (generated in Engelhardt et al. (2019)) using Gateway® LR-cloning to create pGY1-*mCherry-RIC171*. For transient transformation of barley epidermal cells, 1 µg of pGY1-*mCherry-RIC171* was mixed with 1 µg of pGY1-*CFP* and either 0.5 µg of pGY1-*GFP*, 1 µg of pGY1-*GFP*-CA-*RACB* or 1 µg of pGY1-*GFP*-CA-*RACB*-K167R for one reaction. Transient transformation of 7 d old wildtype barley plants (cultivar Golden Promise) was carried out as described above. Single epidermal cells were analyzed 24 h after transformation using the Leica TCS SP5. CFP was excited with the 458 nm Argon laser line and detected between 463 nm and 485 nm using a HyD detector, while GFP was excited with the 488 nm Argon laser line and detected between 500 nm and 550 nm using a PMT and mCherry was excited with the 561 nm laser line and detected between 570 nm and 620 nm using another HyD detector. Individual laser excitation strengths and detector sensitivity settings were used for each cell in order to give high quality images. For all cells, sequentially-scanned Z-stacks with 2 µm step size were taken, ranging from the top of the cell until the full nucleus was captured, with the number of Z-steps being unique for each cell. For each combination of constructs, 15 cells were taken per replicate, with 3 biological replicates in total. For better illustration, the brightness of the representative images in this study was enhanced post scanning using the Leica Application Suite X software V3.5.7.23225 (Leica, Wetzlar, Germany).

## Supporting information

assigned within the peptide sequence (S1 Table), both

were found to phosphorylated in those samples (S2 Table).

identified more than 5,000 phosphopeptides from diverse barley proteins (S3 Table), thus

such as three different barley ROPGAPs (S4 Table).

## Acknowledgements

We acknowledge the technical assistance of Cornelia Marthe (IPK Gatersleben) in generating the transgenic barley used in this study and Dalma Ménesi for her help with protein purifications (IPB, BRC Szeged). We are grateful to the TUM plant technology center for support in propagation of transgenic barley plants.

## Supporting information

**S1 Fig.**
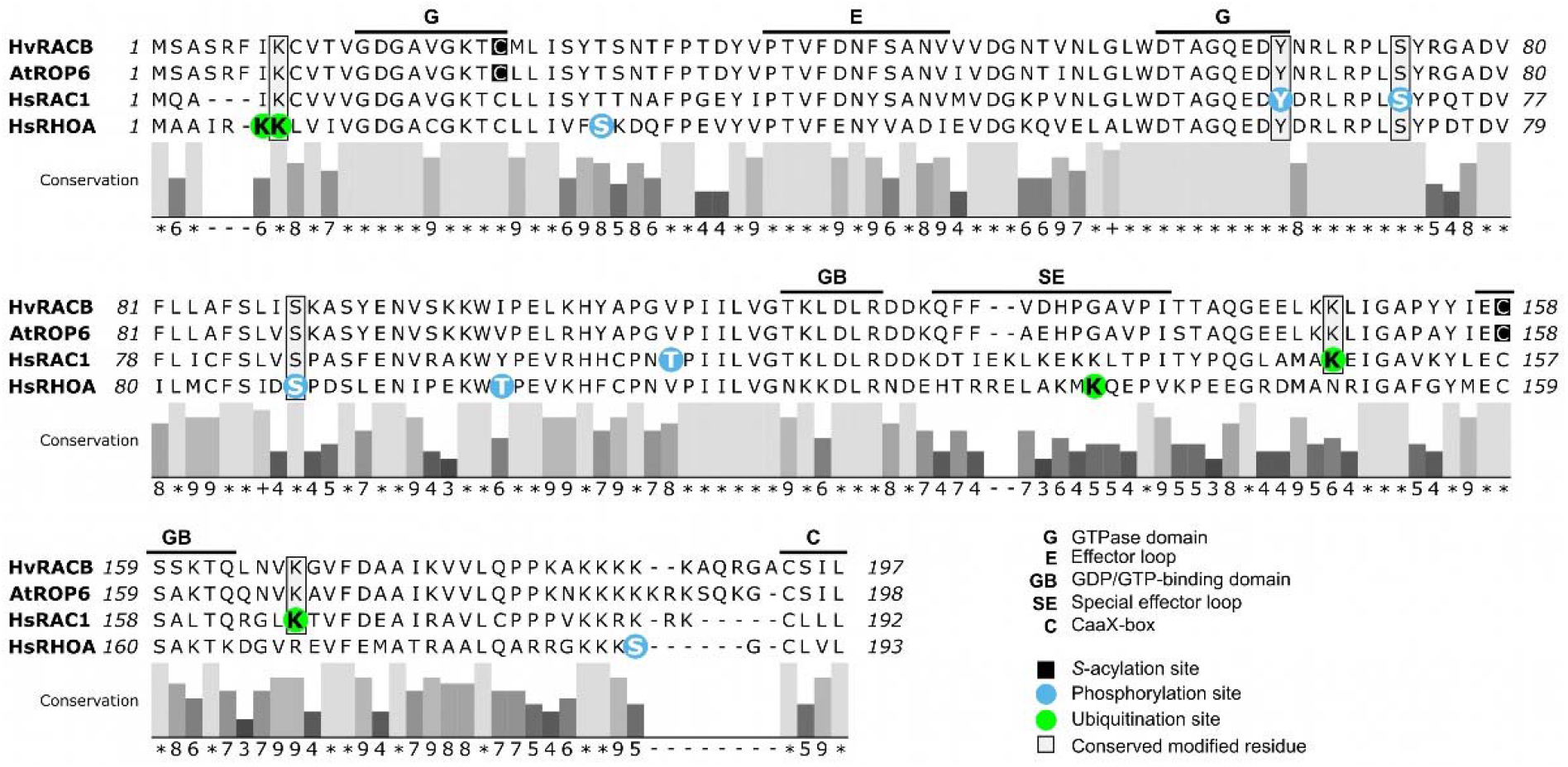
Alignment of RACB with AtROP6, HsRAC1 and HsRHOA. Previously published phosphorylation (blue) and ubiquitination (green) sites of mammalian Rac1 and RhoA are highlighted here in the human RAC1 and RHOA sequences. Conserved sites are highlighted in grey boxes. Additionally, the GTPase domain (G), effector loop (E), nucleotide-binding domain (GB), special effector loop (SE) and CaaX-box prenylation motif (C) are illustrated and were taken from Schultheiss et al. (2003). AtROP6 has been shown to be transiently *S-*acylated at two cysteines (Sorek et al., 2010; Sorek et al., 2017), which are also conserved in RACB (black boxes). Conservation of amino acids was determined using MAFFT multiple-sequence alignment in Jalview (Waterhouse et al., 2009; Katoh and Toh, 2010). Conservation scores range from 0 to 11 and are shown as bars (dark grey = less conserved, light grey = highly conserved). Scores of 0, 10, and 11 are shown as “-“, “+” and “*”, respectively).

**S2 Fig.**
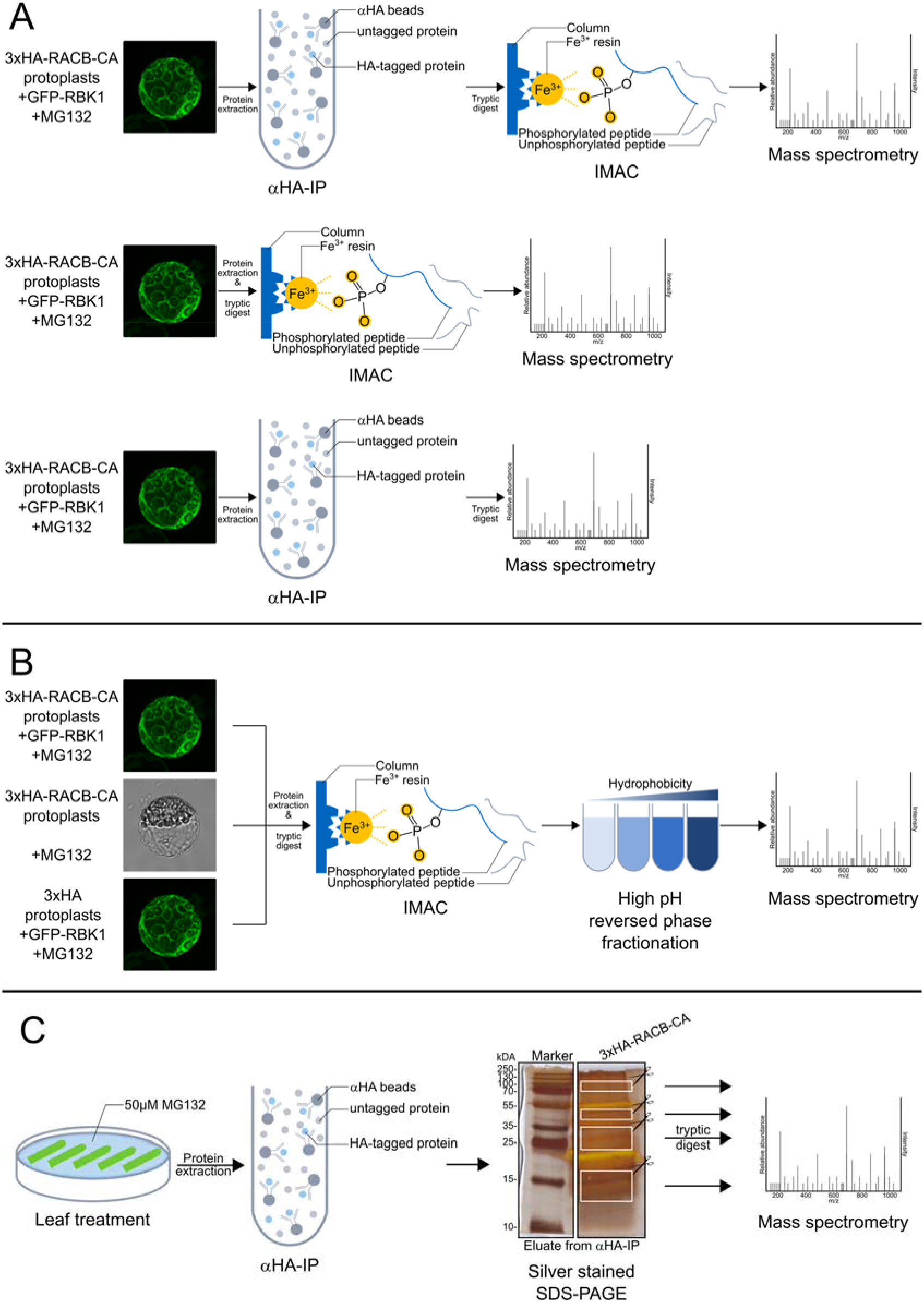
Workflows designed to identify RACB phosphorylation and ubiquitination. **(A)** Identification of phosphorylated peptides after CA-RACB and RBK1 overexpression. Protoplasts were isolated from stably-transgenic barley leaves overexpressing 3xHA-CA-RACB (pooled lines BG657 E3 and E4). These protoplasts were transiently super-transformed with GFP-RBK1 to stimulate phosphorylation events of proteins associated with the RACB/RBK1 pathway. After transformation, all protoplasts were treated with MG132 and proteins were extracted. HA-tagged proteins were first enriched using immunoprecipitation, followed by tryptic digest and phosphopeptide enrichment via IMAC (top panel). In two cases, immunoprecipitation or IMAC were skipped (center and lower panel, respectively). Phosphorylated peptides were identified using mass spectrometry. **(B)** Second experiment to identify phosphorylated peptides after CA-RACB and/or RBK1 overexpression. Protoplasts were isolated from stably transgenic barley leaves overexpressing either 3xHA (lines BG659 E2 and E3) or 3xHA-CA-RACB (lines BG657 E3 and E4). These protoplasts were super-transformed with GFP-RBK1. After transformation, all protoplasts (including an untransformed 3xHA-CA-RACB control) were treated with MG132. Proteins were extracted and digested with trypsin, followed by enrichment of phosphopeptides using IMAC. These samples were decomplexed by fractionating the phosphorylated proteins according to their hydrophobicity. Phosphorylated peptides were identified by mass spectrometry. **(C)** Analysis of CA-RACB ubiquitination by a targeted approach. Transgenic barley leaves overexpressing 3xHA-CA-RACB (from pooled lines BG657 E3 and E4) were floated on a solution containing MG132. HA-containing proteins were enriched by immunoprecipitation and loaded on a SDS-PAGE. Following silver-staining, RACB-containing bands and those corresponding to the higher molecular weight derivatives shown in Fig 5A were excised from the gel (white boxes). After solubilization and tryptic digest, modification of RACB-peptides was investigated via mass spectrometry.

**S3 Fig.**
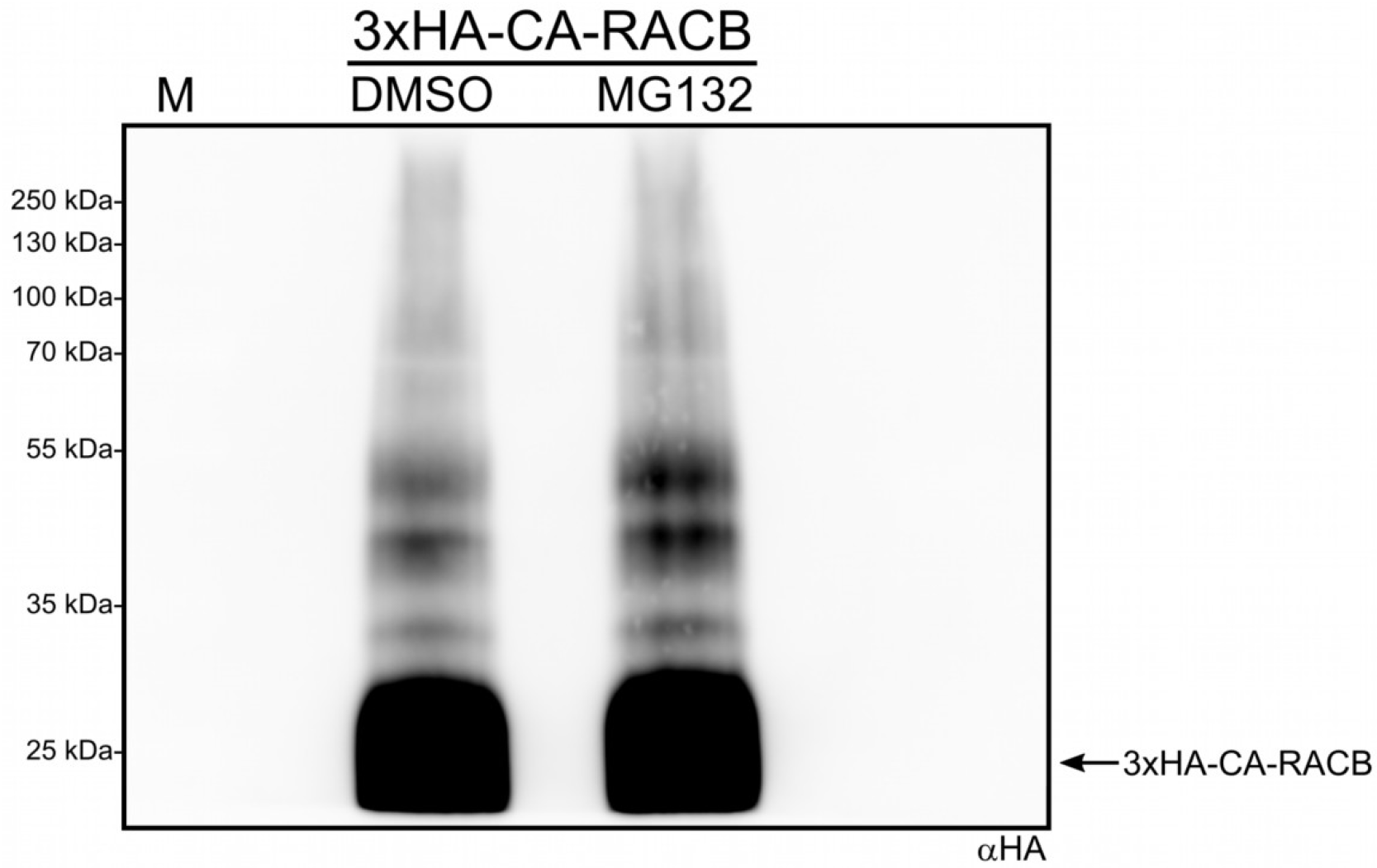
The ubiquitin-like laddering pattern of 3xHA-CA-RACB is independent of MG132. Whole first leaves of barley stably overexpressing 3xHA-CA-RACB were either floated on a solution containing MG132 or DMSO. After protein extraction, αHA-immunoprecipitation and αHA Western blotting, higher molecular weight laddering of 3xHA-CA-RACB could be detected independent of the treatment. Plant material originated from transgenic lines BG657 E3 and E4.

**S4 Fig.**
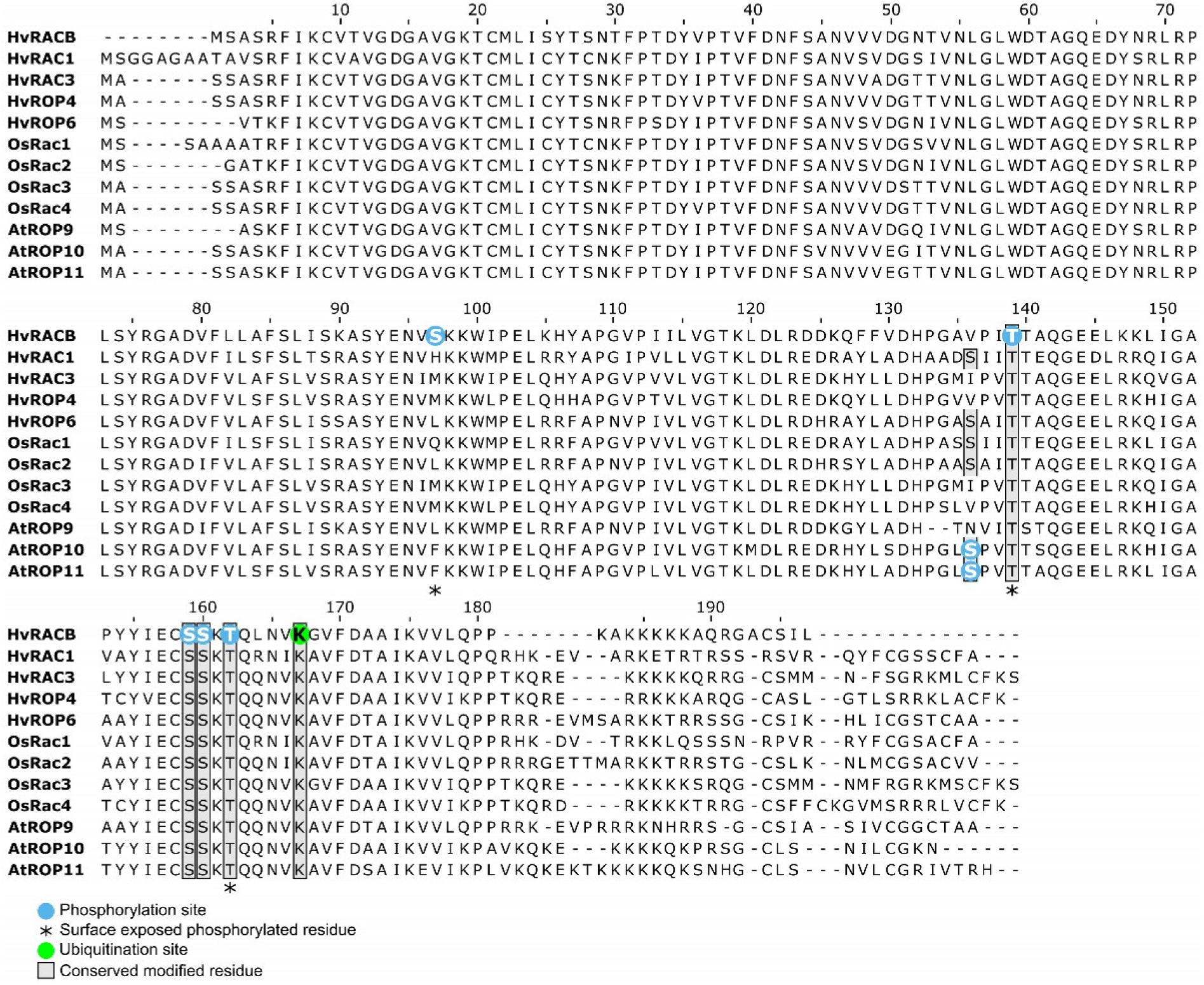
Alignment of barley RACB with type II ROPs from barley, rice *Arabidopsis thaliana*. The CA-RACB ubiquitination site at K167 (green) is conserved in all type II ROPs from the three species (shaded box). *In vitro* phosphorylation sites of RACB (blue) are conserved among most type II ROPs (shaded box). Surface-exposed phosphosites are indicated by asterisks. Conserved amino acids were identified and aligned using MAFFT multiple-sequence alignment in Jalview (Waterhouse et al., 2009; Katoh and Toh, 2010).

**S5 Fig.**
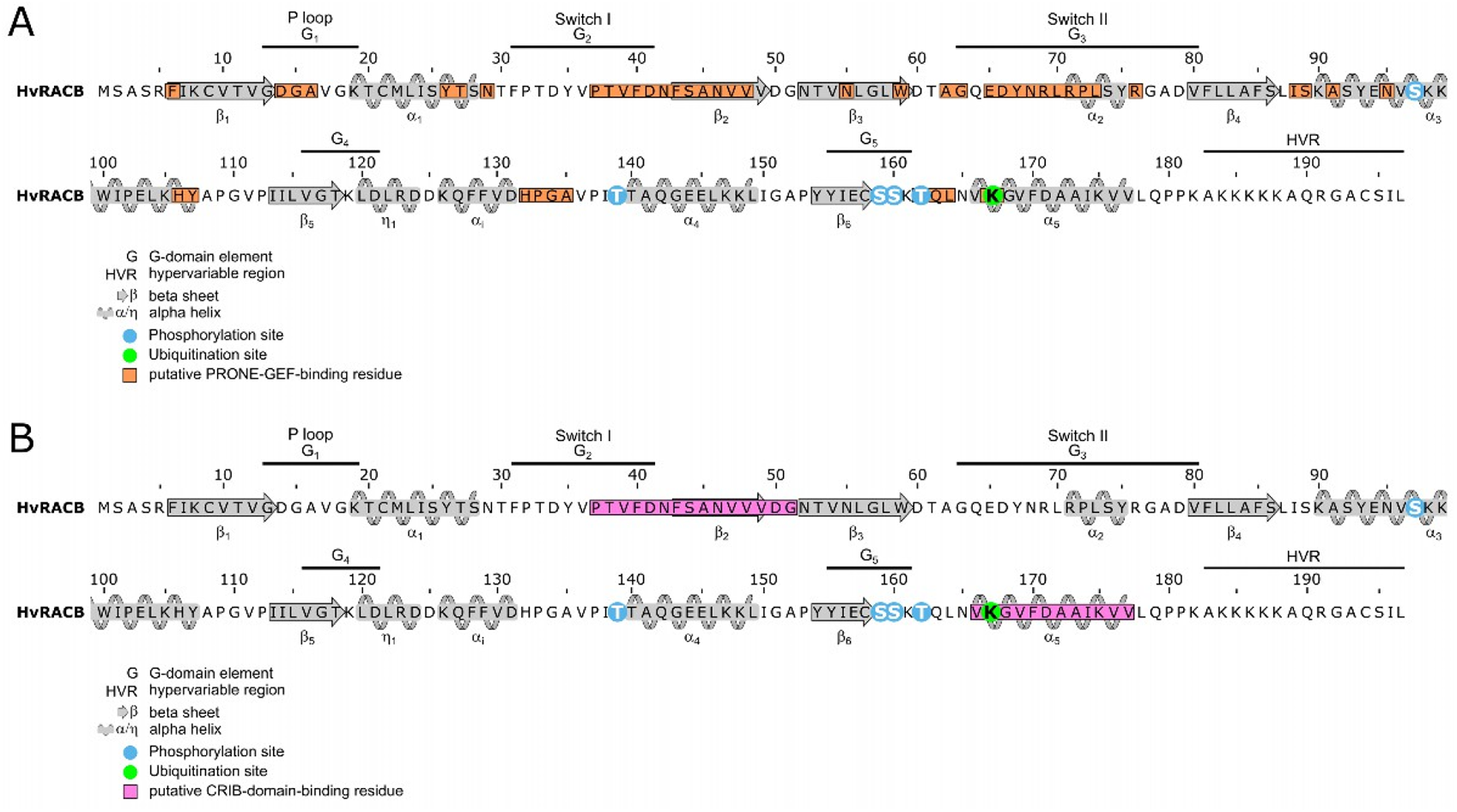
GEF- and CRIB-binding interfaces in RACB. Predicted binding interfaces for PRONE-GEFs (**A**, orange boxes, (Thomas et al., 2007)) and CRIB-domains (**B**, pink boxes, (Abdul-Manan et al., 1999; Schaefer et al., 2011)) are highlighted in the RACB amino acid sequence. RACB secondary structure motifs were obtained through homology-modelling using SWISS-MODEL with the crystal structure of AtROP5 as template (PDB: 3bwd.1, (Fricke and Berken, 2009)). α-helices (grey boxes with winding helices) and β-sheets (grey arrows), as well as RACB *in vitro* phosphorylation (blue circle) and *in vivo* ubiquitination (green circle) sites are illustrated. Annotations for the G-domains (G_1-5_) and hypervariable region (HVR) were taken from Fricke and Berken (2009), whereas regions for P loop and Switches I & II were adopted from Sørmo et al. (2006). Numbering for α-helices and β-sheets follow Pai et al. (1989) and Thomas et al. (2007).

**S1 Table. Dataset of the mass spectrometry analysis of the recombinant RACB and RBK1 *in vitro* kinase assay.**

**S2 Table. Dataset of the first mass-spectrometry experiment to identify *in vivo* phosphorylation of CA-RACB.** This dataset contains all identified barley phosphopeptides originating from transgenic HA-CA-RACB overexpressing protoplasts, which were transformed with GFP-RBK1 and treated with MG132. HA-tagged proteins and/or phosphopeptides were enriched by HA-immunoprecipitation and IMAC, respectively (S2A Fig). All peptides and their modifications were identified by mass spectrometry. Of main interest are the gene identifiers and peptides with phosphorylated residues and associated localization scores.

**S3 Table. Dataset of the second mass-spectrometry experiment to identify *in vivo* phosphorylation of CA-RACB.** Dataset comprising all recovered barley phosphopeptides from protoplasts of stable transgenic barley plants. Phosphopeptides were enriched from full-protein extracts of either HA- or HA-CA-RACB overexpressing protoplasts, which were co-transformed with GFP-RBK1 and incubated with MG132. Samples were decomplexed by fractionating phosphopeptides according to their hydrophobicity (S2B Fig). All peptides and their modifications were identified by mass spectrometry. Non-enriched full-protein samples (FP) were included in mass spectrometry analyses, but did not yield many phosphopeptides. Of main interest in this dataset are the gene identifiers and peptides with phosphorylated amino acids and their corresponding localization scores.

**S4 Table. Summary of all barley ROPGAP phosphorylation sites which were evidenced in this study.**

**S5 Table. List of all primers used in this study.**

## References

Abdrabou, A., and Wang, Z. (2018). Post-translational modification and subcellular distribution of Rac1: An update. Cells 7, 263.

Abdul-Manan, N., Aghazadeh, B., Liu, G.A., Majumdar, A., Ouerfelli, O., Siminovitch, K.A., and Rosen, M.K. (1999). Structure of Cdc42 in complex with the GTPase-binding domain of the ‘Wiskott– Aldrich syndrome’protein. Nature 399, 379–383.

Akamatsu, A., Wong, H.L., Fujiwara, M., Okuda, J., Nishide, K., Uno, K., Imai, K., Umemura, K., Kawasaki, T., and Kawano, Y. (2013). An OsCEBiP/OsCERK1-OsRacGEF1-OsRac1 module is an essential early component of chitin-induced rice immunity. Cell host & microbe 13, 465–476.

Berken, A. (2006). ROPs in the spotlight of plant signal transduction. Cell Mol Life Sci 63, 2446–2459.

Berken, A., and Wittinghofer, A. (2008). Structure and function of Rho-type molecular switches in plants. Plant Physiol Biochem 46, 380–393.

Berken, A., Thomas, C., and Wittinghofer, A. (2005). A new family of RhoGEFs activates the Rop molecular switch in plants. Nature 436, 1176–1180.

Boulter, E., and Garcia-Mata, R. (2010). RhoGDI: A rheostat for the Rho switch. Small GTPases 1, 65–68.

Bradford, M.M. (1976). A rapid and sensitive method for the quantitation of microgram quantities of protein utilizing the principle of protein-dye binding. Anal Biochem 72, 248–254.

Castillo-Lluva, S., Tan, C.T., Daugaard, M., Sorensen, P.H., and Malliri, A. (2013). The tumour suppressor HACE1 controls cell migration by regulating Rac1 degradation. Oncogene 32, 1735–1742.

Chang, F., Lemmon, C., Lietha, D., Eck, M., and Romer, L. (2011). Tyrosine phosphorylation of Rac1: a role in regulation of cell spreading. PLoS One 6, e28587.

Cox, J., and Mann, M. (2008). MaxQuant enables high peptide identification rates, individualized ppb- range mass accuracies and proteome-wide protein quantification. Nature biotechnology 26, 1367–1372.

Dong, S., Zhao, J., Wei, J., Bowser, R.K., Khoo, A., Liu, Z., Luketich, J.D., Pennathur, A., Ma, H., and Zhao, Y. (2014). F-box protein complex FBXL19 regulates TGFbeta1-induced E-cadherin down- regulation by mediating Rac3 ubiquitination and degradation. Mol Cancer 13, 76.

Dorjgotov, D., Jurca, M.E., Fodor-Dunai, C., Szűcs, A., Ötvös, K., Klement, É., Bíró, J., and Fehér, A. (2009). Plant Rho-type (Rop) GTPase-dependent activation of receptor-like cytoplasmic kinases in vitro. FEBS Letters 583, 1175–1182.

Ebine, K., Fujimoto, M., Okatani, Y., Nishiyama, T., Goh, T., Ito, E., Dainobu, T., Nishitani, A., Uemura, T., Sato, M.H., Thordal-Christensen, H., Tsutsumi, N., Nakano, A., and Ueda, T. (2011). A membrane trafficking pathway regulated by the plant-specific RAB GTPase ARA6. Nat Cell Biol 13, 853–859.

Ellerbroek, S.M., Wennerberg, K., and Burridge, K. (2003). Serine phosphorylation negatively regulates RhoA in vivo. J Biol Chem 278, 19023–19031.

Enders, T.A., Frick, E.M., and Strader, L.C. (2017). An Arabidopsis kinase cascade influences auxin- responsive cell expansion. Plant J 92, 68–81.

Engelhardt, S., Kopischke, M., Hofer, J., Probst, K., McCollum, C., and Hückelhoven, R. (2019). Barley RIC157 is involved in RACB-mediated susceptibility to powdery mildew. bioRxiv, 848226.

Engler, C., Kandzia, R., and Marillonnet, S. (2008). A one pot, one step, precision cloning method with high throughput capability. PLoS One 3, e3647.

Fodor-Dunai, C., Fricke, I., Potocky, M., Dorjgotov, D., Domoki, M., Jurca, M.E., Otvos, K., Zarsky, V., Berken, A., and Feher, A. (2011). The phosphomimetic mutation of an evolutionarily conserved serine residue affects the signaling properties of Rho of plants (ROPs). Plant J 66, 669–679.

Forget, M.-A., Desrosiers, R.R., Gingras, D., and Béliveau, R. (2002). Phosphorylation states of Cdc42 and RhoA regulate their interactions with Rho GDP dissociation inhibitor and their extraction from biological membranes. Biochemical Journal 361, 243–254.

Fricke, I., and Berken, A. (2009). Molecular basis for the substrate specificity of plant guanine nucleotide exchange factors for ROP. FEBS letters 583, 75–80.

Gohre, V., Spallek, T., Haweker, H., Mersmann, S., Mentzel, T., Boller, T., de Torres, M., Mansfield, J.W., and Robatzek, S. (2008). Plant pattern-recognition receptor FLS2 is directed for degradation by the bacterial ubiquitin ligase AvrPtoB. Curr Biol 18, 1824–1832.

Hensel, G., Kastner, C., Oleszczuk, S., Riechen, J., and Kumlehn, J. (2009). Agrobacterium-mediated gene transfer to cereal crop plants: current protocols for barley, wheat, triticale, and maize. Int J Plant Genomics 2009, 835608.

Hoefle, C., Huesmann, C., Schultheiss, H., Börnke, F., Hensel, G., Kumlehn, J., and Hückelhoven, R. (2011). A barley ROP GTPase ACTIVATING PROTEIN associates with microtubules and regulates entry of the barley powdery mildew fungus into leaf epidermal cells. The Plant Cell 23, 2422–2439.

Hua, Z., and Vierstra, R.D. (2011). The cullin-RING ubiquitin-protein ligases. Annu Rev Plant Biol 62, 299–334.

Huesmann, C., Reiner, T., Hoefle, C., Preuss, J., Jurca, M.E., Domoki, M., Feher, A., and Huckelhoven, R. (2012). Barley ROP binding kinase1 is involved in microtubule organization and in basal penetration resistance to the barley powdery mildew fungus. Plant Physiol 159, 311–320.

Irani, K., Xia, Y., Zweier, J.L., Sollott, S.J., Der, C.J., Fearon, E.R., Sundaresan, M., Finkel, T., and Goldschmidt-Clermont, P.J. (1997). Mitogenic signaling mediated by oxidants in Ras-transformed fibroblasts. Science 275, 1649–1652.

Jank, T., Bogdanović, X., Wirth, C., Haaf, E., Spoerner, M., Böhmer, K.E., Steinemann, M., Orth, J.H.C., Kalbitzer, H.R., Warscheid, B., Hunte, C., and Aktories, K. (2013). A bacterial toxin catalyzing tyrosine glycosylation of Rho and deamidation of Gq and Gi proteins. Nature Structural & Molecular Biology 20, 1273–1280.

Jones, M.A., Shen, J.-J., Fu, Y., Li, H., Yang, Z., and Grierson, C.S. (2002). The Arabidopsis Rop2 GTPase is a positive regulator of both root hair initiation and tip growth. The Plant Cell 14, 763–776.

Jurca, M.E., Bottka, S., and Fehér, A. (2008). Characterization of a family of Arabidopsis receptor-like cytoplasmic kinases (RLCK class VI). Plant Cell Reports 27, 739–748.

Katoh, K., and Toh, H. (2010). Parallelization of the MAFFT multiple sequence alignment program. Bioinformatics 26, 1899–1900.

Klahre, U., and Kost, B. (2006). Tobacco RhoGTPase ACTIVATING PROTEIN1 Spatially Restricts Signaling of RAC/Rop to the Apex of Pollen Tubes. The Plant Cell 18, 3033–3046.

Krauthammer, M., Kong, Y., Ha, B.H., Evans, P., Bacchiocchi, A., McCusker, J.P., Cheng, E., Davis, M.J., Goh, G., Choi, M., Ariyan, S., Narayan, D., Dutton-Regester, K., Capatana, A., Holman, E.C., Bosenberg, M., Sznol, M., Kluger, H.M., Brash, D.E., Stern, D.F., Materin, M.A., Lo, R.S., Mane, S., Ma, S., Kidd, K.K., Hayward, N.K., Lifton, R.P., Schlessinger, J., Boggon, T.J., and Halaban, R. (2012). Exome sequencing identifies recurrent somatic RAC1 mutations in melanoma. Nat Genet 44, 1006–1014.

Lang, P., Gesbert, F., Delespine-Carmagnat, M., Stancou, R., Pouchelet, M., and Bertoglio, J. (1996). Protein kinase A phosphorylation of RhoA mediates the morphological and functional effects of cyclic AMP in cytotoxic lymphocytes. EMBO J 15, 510–519.

Lawson, C.D., and Ridley, A.J. (2018). Rho GTPase signaling complexes in cell migration and invasion. J Cell Biol 217, 447–457.

Lazo, G.R., Stein, P.A., and Ludwig, R.A. (1991). A DNA transformation-competent Arabidopsis genomic library in Agrobacterium. Biotechnology (N Y) 9, 963–967.

Lin, Y., and Yang, Z. (1997). Inhibition of Pollen Tube Elongation by Microinjected Anti-Rop1Ps Antibodies Suggests a Crucial Role for Rho-Type GTPases in the Control of Tip Growth. Plant Cell 9, 1647–1659.

Lu, D., Lin, W., Gao, X., Wu, S., Cheng, C., Avila, J., Heese, A., Devarenne, T.P., He, P., and Shan, L. (2011). Direct ubiquitination of pattern recognition receptor FLS2 attenuates plant innate immunity. Science 332, 1439–1442.

MacLean, B., Tomazela, D.M., Shulman, N., Chambers, M., Finney, G.L., Frewen, B., Kern, R., Tabb, D.L., Liebler, D.C., and MacCoss, M.J. (2010). Skyline: an open source document editor for creating and analyzing targeted proteomics experiments. Bioinformatics 26, 966–968.

Marx, H., Lemeer, S., Schliep, J.E., Matheron, L., Mohammed, S., Cox, J., Mann, M., Heck, A.J., and Kuster, B. (2013). A large synthetic peptide and phosphopeptide reference library for mass spectrometry–based proteomics. Nature biotechnology 31, 557–564.

McCollum, C., Engelhardt, S., Weiss, L., and Hückelhoven, R. (2020). ROP INTERACTIVE PARTNER b Interacts with RACB and Supports Fungal Penetration into Barley Epidermal Cells. Plant Physiology 184, 823–836.

Mergner, J., Frejno, M., List, M., Papacek, M., Chen, X., Chaudhary, A., Samaras, P., Richter, S., Shikata, H., Messerer, M., Lang, D., Altmann, S., Cyprys, P., Zolg, D.P., Mathieson, T., Bantscheff, M., Hazarika, R.R., Schmidt, T., Dawid, C., Dunkel, A., Hofmann, T., Sprunck, S., Falter-Braun, P., Johannes, F., Mayer, K.F.X., Jürgens, G., Wilhelm, M., Baumbach, J., Grill, E., Schneitz, K., Schwechheimer, C., and Kuster, B. (2020). Mass-spectrometry-based draft of the Arabidopsis proteome. Nature.

Milburn, M.V., Tong, L., deVos, A.M., Brunger, A., Yamaizumi, Z., Nishimura, S., and Kim, S.H. (1990). Molecular switch for signal transduction: structural differences between active and inactive forms of protooncogenic ras proteins. Science 247, 939.

Oberoi, T.K., Dogan, T., Hocking, J.C., Scholz, R.-P., Mooz, J., Anderson, C.L., Karreman, C., Meyer zu Heringdorf, D., Schmidt, G., Ruonala, M., Namikawa, K., Harms, G.S., Carpy, A., Macek, B., Köster, R.W., and Rajalingam, K. (2012). IAPs regulate the plasticity of cell migration by directly targeting Rac1 for degradation. The EMBO journal 31, 14–28.

Opalski, K.S., Schultheiss, H., Kogel, K.H., and Huckelhoven, R. (2005). The receptor-like MLO protein and the RAC/ROP family G-protein RACB modulate actin reorganization in barley attacked by the biotrophic powdery mildew fungus Blumeria graminis f.sp. hordei. Plant J 41, 291–303.

Ozdamar, B., Bose, R., Barrios-Rodiles, M., Wang, H.R., Zhang, Y., and Wrana, J.L. (2005). Regulation of the polarity protein Par6 by TGFbeta receptors controls epithelial cell plasticity. Science 307, 1603–1609.

Pai, E.F., Kabsch, W., Krengel, U., Holmes, K.C., John, J., and Wittinghofer, A. (1989). Structure of the guanine-nucleotide-binding domain of the Ha-ras oncogene product p21 in the triphosphate conformation. Nature 341, 209–214.

Pathuri, I.P., Zellerhoff, N., Schaffrath, U., Hensel, G., Kumlehn, J., Kogel, K.H., Eichmann, R., and Huckelhoven, R. (2008). Constitutively activated barley ROPs modulate epidermal cell size, defense reactions and interactions with fungal leaf pathogens. Plant Cell Rep 27, 1877–1887.

Peng, J., Schwartz, D., Elias, J.E., Thoreen, C.C., Cheng, D., Marsischky, G., Roelofs, J., Finley, D., and Gygi, S.P. (2003). A proteomics approach to understanding protein ubiquitination. Nat Biotechnol 21, 921–926.

Rademacher, E.H., and Offringa, R. (2012). Evolutionary Adaptations of Plant AGC Kinases: From Light Signaling to Cell Polarity Regulation. Front Plant Sci 3, 250.

Reiner, T., Hoefle, C., and Hückelhoven, R. (2016). A barley SKP1-like protein controls abundance of the susceptibility factor RACB and influences the interaction of barley with the barley powdery mildew fungus. Molecular plant pathology 17, 184–195.

Reiner, T., Hoefle, C., Huesmann, C., Ménesi, D., Fehér, A., and Hückelhoven, R. (2015). The Arabidopsis ROP-activated receptor-like cytoplasmic kinase RLCK VI_A3 is involved in control of basal resistance to powdery mildew and trichome branching. Plant Cell Reports 34, 457–468.

Rolli-Derkinderen, M., Sauzeau, V., Boyer, L., Lemichez, E., Baron, C., Henrion, D., Loirand, G., and Pacaud, P. (2005). Phosphorylation of serine 188 protects RhoA from ubiquitin/proteasome-mediated degradation in vascular smooth muscle cells. Circ Res 96, 1152–1160.

Ruprecht, B., Zecha, J., Zolg, D.P., and Kuster, B. (2017a). High pH Reversed-Phase Micro-Columns for Simple, Sensitive, and Efficient Fractionation of Proteome and (TMT labeled) Phosphoproteome Digests. Methods Mol Biol 1550, 83–98.

Ruprecht, B., Koch, H., Domasinska, P., Frejno, M., Kuster, B., and Lemeer, S. (2017b). Optimized Enrichment of Phosphoproteomes by Fe-IMAC Column Chromatography. Methods Mol Biol 1550, 47–60.

Schaefer, A., Höhner, K., Berken, A., and Wittinghofer, A. (2011). The unique plant RhoGAPs are dimeric and contain a CRIB motif required for affinity and specificity towards cognate small G proteins. Biopolymers 95, 420–433.

Schindelin, J., Arganda-Carreras, I., Frise, E., Kaynig, V., Longair, M., Pietzsch, T., Preibisch, S., Rueden, C., Saalfeld, S., and Schmid, B. (2012). Fiji: an open-source platform for biological-image analysis. Nature methods 9, 676–682.

Schrodinger, LLC. (2015). The PyMOL Molecular Graphics System, Version 1.8.

Schultheiss, H., Dechert, C., Kogel, K.-H., and Hückelhoven, R. (2002). A small GTP-binding host protein is required for entry of powdery mildew fungus into epidermal cells of barley. Plant Physiology 128, 1447–1454.

Schultheiss, H., Dechert, C., Kogel, K.H., and Huckelhoven, R. (2003). Functional analysis of barley RAC/ROP G-protein family members in susceptibility to the powdery mildew fungus. Plant J 36, 589–601.

Schultheiss, H., Preuss, J., Pircher, T., Eichmann, R., and Huckelhoven, R. (2008). Barley RIC171 interacts with RACB in planta and supports entry of the powdery mildew fungus. Cell Microbiol 10, 1815–1826.

Schweizer, P., Pokorny, J., Abderhalden, O., and Dudler, R. (1999). A Transient Assay System for the Functional Assessment of Defense-Related Genes in Wheat. Molecular Plant-Microbe Interactions 12, 647–654.

Sorek, N., Poraty, L., Sternberg, H., Buriakovsky, E., Bar, E., Lewinsohn, E., and Yalovsky, S. (2017). Corrected and Republished from: Activation Status-Coupled Transient S-Acylation Determines Membrane Partitioning of a Plant Rho-Related GTPase. Molecular and Cellular Biology 37, e00333–00317.

Sorek, N., Segev, O., Gutman, O., Bar, E., Richter, S., Poraty, L., Hirsch, J.A., Henis, Y.I., Lewinsohn, E., Jürgens, G., and Yalovsky, S. (2010). An S-Acylation Switch of Conserved G Domain Cysteines Is Required for Polarity Signaling by ROP GTPases. Current Biology 20, 914–920.

Sorek, N., Gutman, O., Bar, E., Abu-Abied, M., Feng, X., Running, M.P., Lewinsohn, E., Ori, N., Sadot, E., Henis, Y.I., and Yalovsky, S. (2011). Differential effects of prenylation and s-acylation on type I and II ROPS membrane interaction and function. Plant Physiol 155, 706–720.

Sørmo, C.G., Leiros, I., Brembu, T., Winge, P., Os, V., and Bones, A.M. (2006). The crystal structure of Arabidopsis thaliana RAC7/ROP9: the first RAS superfamily GTPase from the plant kingdom. Phytochemistry 67, 2332–2340.

Stanislas, T., Huser, A., Barbosa, I.C., Kiefer, C.S., Brackmann, K., Pietra, S., Gustavsson, A., Zourelidou, M., Schwechheimer, C., and Grebe, M. (2015). Arabidopsis D6PK is a lipid domain-dependent mediator of root epidermal planar polarity. Nat Plants 1, 15162.

Studer, G., Rempfer, C., Waterhouse, A.M., Gumienny, R., Haas, J., and Schwede, T. (2020). QMEANDisCo-distance constraints applied on model quality estimation. Bioinformatics 36, 1765–1771.

Tang, J., Ip, J.P.K., Ye, T., Ng, Y.-P., Yung, W.-H., Wu, Z., Fang, W., Fu, A.K.Y., and Ip, N.Y. (2014). Cdk5- Dependent Mst3 Phosphorylation and Activity Regulate Neuronal Migration through RhoA Inhibition. The Journal of Neuroscience 34, 7425–7436.

Thomas, C., Fricke, I., Scrima, A., Berken, A., and Wittinghofer, A. (2007). Structural evidence for a common intermediate in small G protein-GEF reactions. Mol Cell 25, 141–149.

Tong, J., Li, L., Ballermann, B., and Wang, Z. (2016). Phosphorylation and Activation of RhoA by ERK in Response to Epidermal Growth Factor Stimulation. PLoS One 11, e0147103.

Torrino, S., Visvikis, O., Doye, A., Boyer, L., Stefani, C., Munro, P., Bertoglio, J., Gacon, G., Mettouchi, A., and Lemichez, E. (2011). The E3 ubiquitin-ligase HACE1 catalyzes the ubiquitylation of active Rac1. Dev Cell 21, 959–965.

Vickers, C.E., Xue, G.P., and Gresshoff, P.M. (2003). A synthetic xylanase as a novel reporter in plants. Plant Cell Reports 22, 135–140.

Vizcaíno, J.A., Csordas, A., Del-Toro, N., Dianes, J.A., Griss, J., Lavidas, I., Mayer, G., Perez-Riverol, Y., Reisinger, F., and Ternent, T. (2016). 2016 update of the PRIDE database and its related tools. Nucleic acids research 44, D447–D456.

Wang, H.-R., Zhang, Y., Ozdamar, B., Ogunjimi, A.A., Alexandrova, E., Thomsen, G.H., and Wrana, J.L. (2003). Regulation of Cell Polarity and Protrusion Formation by Targeting RhoA for Degradation. Science 302, 1775–1779.

Waterhouse, A., Bertoni, M., Bienert, S., Studer, G., Tauriello, G., Gumienny, R., Heer, F.T., de Beer, T.A.P., Rempfer, C., and Bordoli, L. (2018). SWISS-MODEL: homology modelling of protein structures and complexes. Nucleic acids research 46, W296–W303.

Waterhouse, A.M., Procter, J.B., Martin, D.M., Clamp, M., and Barton, G.J. (2009). Jalview Version 2--a multiple sequence alignment editor and analysis workbench. Bioinformatics 25, 1189–1191.

Wei, J., Mialki, R.K., Dong, S., Khoo, A., Mallampalli, R.K., Zhao, Y., and Zhao, J. (2013). A new mechanism of RhoA ubiquitination and degradation: roles of SCF(FBXL19) E3 ligase and Erk2. Biochim Biophys Acta 1833, 2757–2764.

Wennerberg, K., Rossman, K.L., and Der, C.J. (2005). The Ras superfamily at a glance. J Cell Sci 118, 843–846.

Worby, C.A., Mattoo, S., Kruger, R.P., Corbeil, L.B., Koller, A., Mendez, J.C., Zekarias, B., Lazar, C., and Dixon, J.E. (2009). The fic domain: regulation of cell signaling by adenylylation. Mol Cell 34, 93–103.

Wu, G., Li, H., and Yang, Z. (2000). Arabidopsis RopGAPs Are a Novel Family of Rho GTPase-Activating Proteins that Require the Cdc42/Rac-Interactive Binding Motif for Rop-Specific GTPase Stimulation. Plant Physiology 124, 1625–1636.

Yalovsky, S. (2015). Protein lipid modifications and the regulation of ROP GTPase function. J Exp Bot 66, 1617–1624.

Yang, Z. (2008). Cell polarity signaling in Arabidopsis. Annu Rev Cell Dev Biol 24, 551–575.

Zhao, J., Mialki, R.K., Wei, J., Coon, T.A., Zou, C., Chen, B.B., Mallampalli, R.K., and Zhao, Y. (2013). SCF E3 ligase F-box protein complex SCFFBXL19 regulates cell migration by mediating Rac1 ubiquitination and degradation. The FASEB Journal 27, 2611–2619.

Zolg, D.P., Wilhelm, M., Yu, P., Knaute, T., Zerweck, J., Wenschuh, H., Reimer, U., Schnatbaum, K., and Kuster, B. (2017). PROCAL: A Set of 40 Peptide Standards for Retention Time Indexing, Column Performance Monitoring, and Collision Energy Calibration. PROTEOMICS 17, 1700263.

Zolg, D.P., Wilhelm, M., Schmidt, T., Medard, G., Zerweck, J., Knaute, T., Wenschuh, H., Reimer, U., Schnatbaum, K., and Kuster, B. (2018). ProteomeTools: Systematic characterization of 21 post-translational protein modifications by liquid chromatography tandem mass spectrometry (LC-MS/MS) using synthetic peptides. Molecular & Cellular Proteomics 17, 1850–1863.

